# Maturation and Refinement of the Maculae and Foveae in the Anolis sagrei lizard

**DOI:** 10.1101/2023.02.18.529081

**Authors:** M. Austin Wahle, Hannah Q. Kim, Douglas B. Menke, James D. Lauderdale, Ashley M. Rasys

## Abstract

In primates, the fovea is a pit in the center of the macula, which is a region of the retina with a high concentration of cone photoreceptor cells that accounts for a large degree of visual acuity in primates. The maturation of this primate visual acuity area is characterized by the shallowing and widening of the foveal pit, a decrease in the diameter of the rod-free zone, and packing of photoreceptor cells more densely into this area of the retina, which occurs sometime after birth. Maturation occurs concurrently with progressing age and development, which further correlates with increasing eye size, retinal length, and retinal area. These observations have led to the hypothesis that the maturation of the fovea might be a function of mechanical variables that remodel the retina. However, this has never been explored outside of primates. Here, we take advantage of the *Anolis sagrei* lizard, which has a bifoveated retina, to study maturation of the fovea and macula. Eyes were collected from male and female lizards from five different age groups – hatchling, 2-month, 4-month, 6-month, and adult. We found that *Anolis* maculae undergo a maturation process somewhat different than what has been observed in primates. Anole macular diameters actually increase in size and undergo minimal photoreceptor cell packing, possessing a near complete complement of these cells at the time of hatching. As the anole eye expands, foveal centers experience little change in overall retina cell density with most cell redistribution occurring at the macular borders and peripheral retina areas.

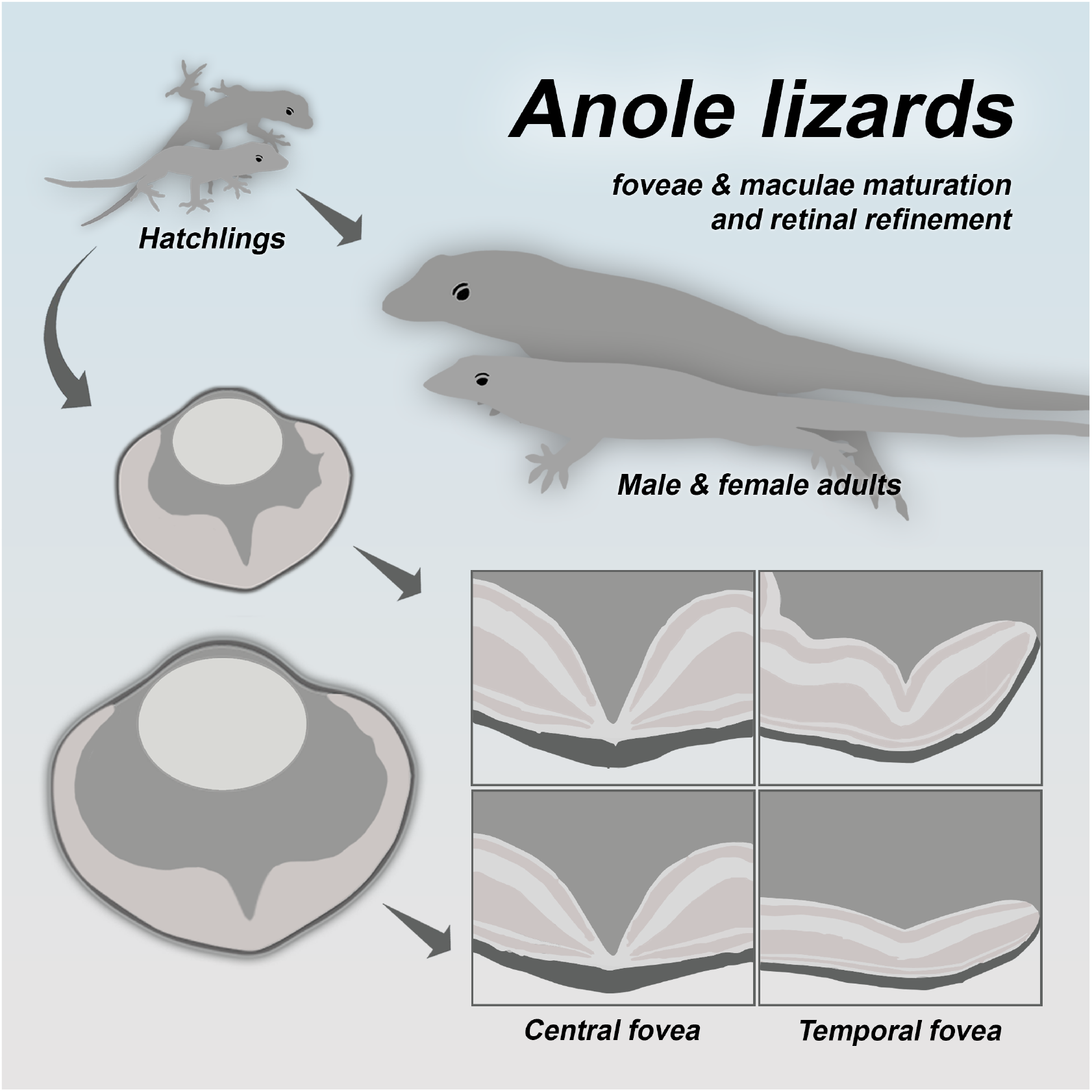

**Highlights:**

- *Foveal and retinal refinement occurs post-hatching into adulthood in anoles*.
- *The temporal pit shallows and widens, whereas central pit morphology remains relatively constant with age*.
- *Retina cell number within a section decreases as the eye grows larger with age*.
- *Anole macular diameter increases from hatching to adult but does not increase proportionately to overall eye growth*.

## 1. Introduction

Since its discovery in the 1800s, the fovea, which is a pit in the back of the retina of the eye, has remained a topic of interest for scientists and philosophers alike because of its importance for visual acuity (Slonaker, 1897). Although the eyes of humans, other haplorrhine primates, some birds, lizards, and fish contain these enigmatic structures (Easter, 1992; Fite & Lister, 1981; Fite & Rosenfield-Wessels, 1975; Mann, 1964; Slonaker, 1897; Walls, 1942), our knowledge of this structure primarily comes from studies performed in primates, with more limited contributions coming from studies of lizards and birds (Potier et al., 2020; Rasys, Pau, Irwin, Luo, Kim, Wahle, Trainor, et al., 2021; Springer & Hendrickson, 2004a, 2004b, 2005; Sugiyama et al., 2020).

The vertebrate retina is typically composed of three cell layers – the ganglion, inner nuclear, and outer nuclear cell layers. The outer nuclear layer contains both rod and cone photoreceptor cells. Cones are responsible for processing high-light stimuli, and rods process low-light stimuli and are absent from the mature central region of the macula (Provis & Hendrickson, 2008; Yuodelis & Hendrickson, 1986). During late embryonic development in primates, the foveal pit is formed by the outward displacement of the ganglion and inner nuclear cell layers. Cone photoreceptor cells within the outer nuclear layer then pack in around the foveal center, which initiates the formation of a focal area of high visual acuity within the retina – the macula. After birth, this area of visual acuity continues to mature by the shallowing and widening of the foveal pit and increasing macular cone photoreceptor cell density (Hendrickson et al., 2012b; Springer & Hendrickson, 2005).

The packing of cones leads to the shrinking of the diameter of the retina’s pure cone ‘rod-free zone’. This process takes around 15 months after birth for the pit to reach maturity and up to 4yrs for the macula to reach peak cone cell volume and a consistent rod free zone diameter (Hendrickson et al., 2012a; Yuodelis & Hendrickson, 1986). Studies investigating this have also found that maturation of the macular region correlates with periods of ocular growth and the expansion of the retina after birth (Springer & Hendrickson, 2004b, 2005). This has led to the hypothesis that the remodeling of the fovea and the centripetal movement of photoreceptor cells might be a function of mechanical variables, namely intraocular pressure and forces generated through ocular-growth-induced retina stretch (Packer et al., 1990; Provis et al., 2013; Schmid et al., 2003; Springer & Hendrickson, 2004a, 2004b, 2005). However, this remains to be functionally tested in primates or investigated in other foveated animals.

Outside of primates and birds, the only other foveated vertebrate to have the development of its eye studied is the brown anole lizard (*Anolis sagrei*) (Rasys, Pau, Irwin, Luo, Kim, Wahle, Trainor, et al., 2021; Rasys, Pau, Irwin, Luo, Kim, Wahle, Menke, et al., 2021; Rasys, Pau, Irwin, Luo, Menke, et al., 2021). This species relies on sight as its principal sense and requires a high degree of visual acuity for hunting its prey and communicating and socially interacting with others of its kind (Fleishman et al., 1995; Fleishman & Pallus, 2010). The brown anole retina is adapted for high-acuity diurnal vision and develops two distinct foveae in each eye during embryogenesis, one located in the center of the retina and one oriented in the temporal region (Fleishman, 1992; Makaretz & Levine, 1980; Rasys, Pau, Irwin, Luo, Kim, Wahle, Trainor, et al., 2021; Rasys, Pau, Irwin, Luo, Kim, Wahle, Menke, et al., 2021). The central fovea is deep and convexiclivate in morphology while the temporal fovea is shallower and more variable in shape and retinal location across different anole species (Fite & Lister, 1981; Fite & Rosenfield-Wessels, 1975).

During late embryonic development, *Anolis* foveae form through the remodeling of the retina cell layers, much like in primates (Hendrickson et al., 2012b; Springer & Hendrickson, 2004b). All of the retinal cell layers become laterally displaced within the central foveal region shortly before hatching. However, the temporal foveal region retains these layers during the formation of the pit. Photoreceptor cell packing in the foveal areas also occurs during this period, and by the time of hatching, the anole can hunt (Rasys, Pau, Irwin, Luo, Kim, Wahle, Trainor, et al., 2021; Rasys, Pau, Irwin, Luo, Kim, Wahle, Menke, et al., 2021). Although this might suggest a high degree of visual acuity, it is unclear whether photoreceptor numbers in the foveal region continue to increase during post-hatchling periods as the eye continues to expand. The retinal landscape does appear to undergo remodeling post-hatching, as can be seen by the shallowing and widening of the temporal fovea (Fig. 1). This observation prompted us to speculate that the post-hatching, anoles may undergo a foveal maturation process analogous to the process that occurs in early postnatal primates.

**Figure 1.**
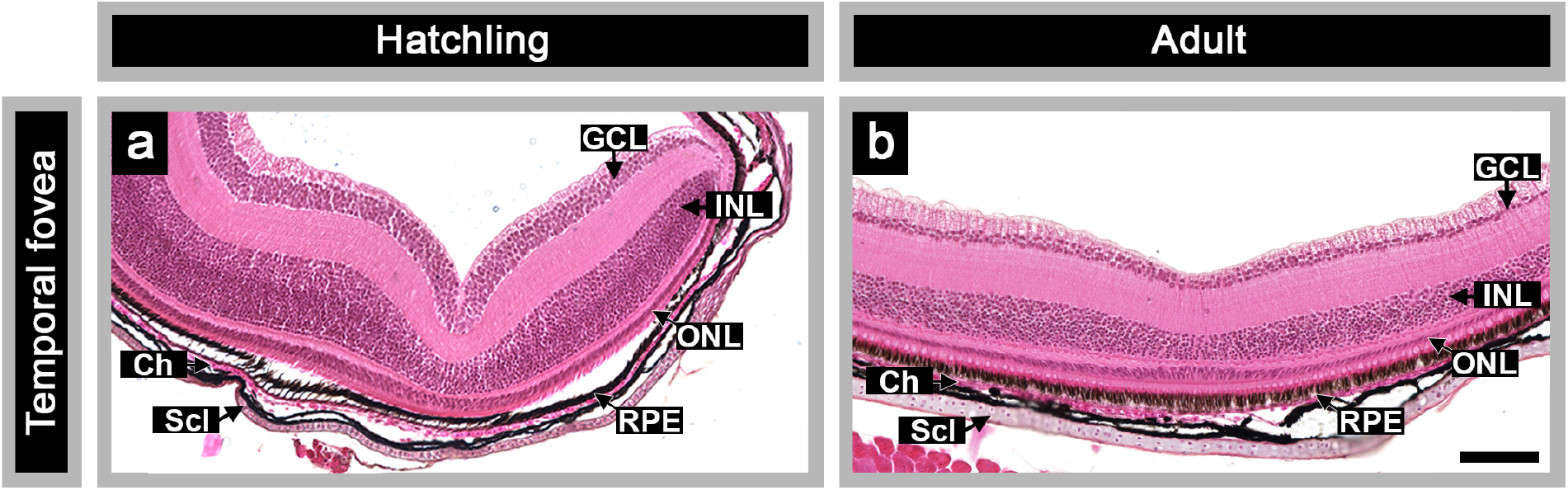
Hematoxylin and eosin stained retinal sections showing the temporal fovea of (A) a hatchling lizard and (B) an adult lizard. The scalebar represents 50 microns. The GCL (ganglion cell layer), INL (inner nuclear layer), and ONL (outer nuclear layer) are denoted along with the RPE (retinal pigmented epithelium), Ch (choroid), and Scl (sclera).

To investigate *Anolis* foveal maturation, we examined the eyes of lizards at five different time points post-hatching: hatchling (<1 week), 2-month, 4-month, 6-month, and adult (>1 year). We evaluated growth trends for overall body size and eye size and then serially sectioned the eyes to relate these findings to gross morphology of the retina, retinal length, retinal depth, pit morphology, global retinal cell populations, and macular cell populations. Through these analyses, we characterized the maturation process of the brown anole retina.

## 2. Methods

### 2.1 Animals

*Anolis sagrei* lizards were housed and bred in a colony at the University of Georgia following the guidelines by Sanger, et al., 2008 (Sanger et al., 2008). Eggs were collected weekly and incubated in vermiculite at 27.8°C and 70% humidity. Upon hatching, lizards were raised until 2-, 4-, and 6-months of age. Wild caught adult lizards that were maintained in the colony for at least 1 year were also included in this study. A minimum of ten lizards were collected for each sex within each age group. This experiment was conducted in accordance with the National Institutes of Health Guide for the Care and Use of Laboratory Animals and was performed with approval and oversight of the University of Georgia Institutional Animal Care and Use Committee.

### 2.2 Euthanasia

Lizards in the designated age groups were first anesthetized with an injection of 250mg/kg of Tricaine (MS-222; Western Chemical Inc) into the coelomic cavity. After loss of consciousness and any response to noxious stimuli, a second lethal injection of MS-222 was made directly into the heart. Lizards then underwent cervical dislocation. Prior to euthanasia, body masses were recorded. Following euthanasia, snout to vent measurements were documented and dissections were performed to identify and

confirm sex. Sex was also independently verified with PCR genotyping. DNA was extracted from the livers of each lizard using standard protocols. PCR was performed using the following primers:

Krank1 AcF 5’-CCTTCCTTTGTAGGATCCAGT-3’
Krank1 AcR 5’-GGAGCACAGGGATAGTTTTGAC-3’
AsagB-F1 5’-GAAGACCAGGAGAGCAARGTC-3’
AsagB-R1 5’-GATGTCGGCAGCYTTGCGTAC-3’

### 2.3 Dissection & Imaging

Eyes were removed from skulls using blunt forceps, placed into 1x phosphate-buffered saline (PBS; 137 mM NaCl, 2.7 mM KCl, 10 mM 16 Na2HPO4, 1.8 mM KH2PO4, pH 7.4), and then imaged with a ZEISS Discovery V12 SteREO microscope and AxioCam camera. Dorsal and lateral views of each eye were taken and measurements along the nasal-temporal axis (x-axis), dorsal-ventral axis (y-axis), and the depth of the eye (z-axis) were made for a minimum of ten lizards for each sex within each age group. These axial measurements were then used to calculate an overall ocular dimension by taking the mean of all three axes. Eyes were then fixed with Bouin’s Solution overnight at 4°C on a rocker.

### 2.4 Tissue Processing and Paraffin Sectioning

Eyes were processed using PBS, ethanol, and Xylene solutions along with paraffin waxes as previously reported (Rasys, Pau, Irwin, Luo, Kim, Wahle, Trainor, et al., 2021). Eyes were then serial sectioned in paraffin wax on the horizontal plane along the dorsal-ventral axis. Sections were then stained with hematoxylin & eosin and mounted in Cytoseal (Thermo Scientific™ Richard-Allan Scientific™) using a protocol previously described in *Rasys et al., 2021* (Rasys, Pau, Irwin, Luo, Kim, Wahle, Trainor, et al., 2021).

### 2.5 Criteria for a well-sectioned eye

An eye was considered perfectly sectioned if retina center, lens center, central fovea, and temporal fovea were within 5 optical degrees of each other. ‘Retina center’ sections were identified by counting the total number of sections containing retina and taking the middlemost section. In a similar manner, the section containing ‘lens center’ was also identified by tallying up the total number of sections containing a lens and then identifying the middlemost section. The central fovea section was determined by counting the total number of sections where retinal cell layers were absent within the pit center and selecting the middle-most section. This was not possible to do with the temporal fovea because all the retinal cell layers are retained within its pit. Instead, temporal fovea sections were identified by selecting the section with the deepest pit. This selection was validated by counting the total number of sections with a pit and identifying the middlemost section.

### 2.6 Imaging & retinal binning

After ensuring that eyes met the above criteria, retina sections with central and temporal foveae were then imaged at both 20x and 40x magnification using the KEYENCE BZ-700 microscope and Keyence stitching software program to generate a finalized set of images for performing cell counts. We binned the 20x and 40x images into segments that would allow for comparison of the eyes across age and sex. Binning the eyes addressed the unique geometry of the brown anole eye, ensuring that the central fovea pit, temporal fovea pit, parafoveal regions, and peripheral retina could be meaningfully compared across age and sex.

The 20x images captured the entire retina and were used to obtain total retina cell counts from a section. The retina was first segmented into ten bins by measuring the inner and outer total retinal lengths using Adobe Illustrator (Supp. Fig. 1a). An important consideration before binning was that the location of the central fovea is biased towards the temporal side of the retina, and not aligned with the central axis of the eye. Therefore, we sought to distinguish the different axes within the eye. The central axis is represented by the z-axis (depth) of the eye and spans the distance between the cornea and the central retina, effectively dividing the eye into two equal halves. We defined the visual axes as being the dimension from the cornea to the foveal centers, which was measured to be 5° temporal of the central axis for the central fovea and 45° for the temporal fovea (Supp. Fig. 1b). The nasal-temporal distance, which represents ocular width, was portrayed as the x-axis. To ensure that sampling of the retinal areas was consistent across age groups with different eye sizes, bins were centered on each of the foveae. As a result, bins N5 and T4 were proportional to each other but slightly reduced compared to the other bins; T2, on the other hand, was slightly larger. This treatment was consistently applied across all retinal sections for every age group to ensure proper retina proportions were retained. This method of binning resulted in five nasal retina bins and four temporal retina bins (Supp. Fig. 1a), which were designated as N5-N1, CF, T1, T2, TF, and T4. When central and temporal foveae were not on the same plane of section, global cell counts from the two separate sections were merged together by using cell counts of bins N5-T1 from the section containing the central fovea, bins TF and T4 from the section with the temporal fovea. We then used the average retinal lengths and cell counts from both sections to generate data for bin T2. This method was selected after it was determined that there was no significant change in the cell number in any of the retinal bins, excluding the bins CF and TF, between the two sections.

The 40x images captured the retina regions immediately surrounding each fovea and were used to analyze the cellular distribution within the foveal regions. For this analysis, binning by optical degrees was preferred over retinal length. Doing so ensured that comparable regions of each macula were analyzed across all age groups and variables like eye size have limited impact on outcomes. We previously found that for both hatchlings and adults, the central macula always fell within a 40 optical degree distance, while the temporal fovea typically spanned a gap less than 20 degrees. Therefore, for the central fovea, we used the central axis (z-axis, the depth of the eye) to calculate the arc length (s=r*θ*) necessary to capture 40° of each lizard’s retina surrounding the central fovea. For the temporal fovea, we used the nasotemporal axis (x-axis, the width of the eye) to capture 20° total surrounding the temporal fovea.

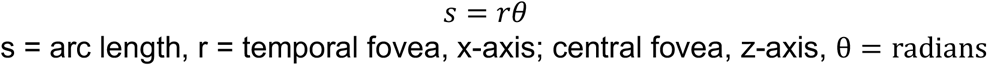

The axial dimensions necessary for these calculations were derived from the measurements made from whole eyes prior to fixing to avoid artifacts that could have occurred during tissue processing and sectioning. Each respective fovea center was then considered the halfway point, and the surrounding retina was then subdivided into 1° bins. This resulted in 40 bins total for the central fovea region and 20 bins total for the temporal fovea (Supp. Fig. 1b). Because different axial measurements were used to calculate central and temporal arc lengths, temporal bins were slightly larger than the central bins.

### 2.7 Cell counts

Automated cell counts were performed using a pipeline that we specifically optimized to handle hematoxylin and eosin-stained retina tissue. This pipeline utilizes multiple Python libraries, including Ski-kit image (van der Walt et al., 2014), NumPy (Harris et al., 2020), matlplotlib (Hunter, 2007), and pandas (McKinney, 2010). In brief, following binning the retinal cell layers (GCL, INL, ONL) were separated using Adobe Photoshop 2021 (version 22.4.2) and saved as individual images. Using the Ski-kit image library (van der Walt et al., 2014), these images were then run through a series of background reduction, color deconvolution, color binarization, and additional noise reduction methods (Ruifrok, 2001; van der Walt et al., 2014) (see GitHub link provided below for specifics).

After filtering, the images underwent Laplacian of Gaussian blot detection for the identification and quantification of cell numbers in each image (Supp. Fig. 1c) (Hongming et al., 2017; Kong et al., 2013). This allowed for the quantification of cells in each retinal cell layer within each bin of each eye. These cell counts were then calculated and recorded using a NumPy dictionary and exported as an excel file into a library dataset using pandas. Matplotlib was used for graphical visualization and error analysis. Accuracy of the pipeline was determined by performing manual cell counts on a random sampling of the datasets (10 bins) as a quality control. The pipeline slightly undercounted the number of cells present compared to manual cell counts (mean accuracy equaled 92.7% ± 4.3%; range 86.8-99.9%). Cell counts were either depicted as raw numbers or as cell densities, which were measured by taking the number of cells divided by the respective bin length.

### 2.7 Defining macular diameter

Maculae were defined as regions of the retina containing a pit along with the adjacent area with increased photoreceptor cell density (i.e., includes area immediately adjacent to the pit, also known as the parafoveal region; Fig. 3). Macular diameters were determined through direct visual assessments as well as through an independent method by setting cellular thresholds.

For visual assessments, we classified the macula boundaries as the area where photoreceptor cell density around the pit tapered to a consistency with the peripheral retina (i.e., the ONL decreases to a single cell layer). This is also where parafoveal thickness decreases. Macula boundaries are more pronounced and easily identified by indentations or shallow depressions present on the retinal surface flanking each of the parafoveal regions in hatchlings and juvenile lizards, but as the eye expands, these areas become more subtle in adults (Fig. 3). Macula diameters determined by visual assessment were made by measuring the horizontal distance between each side of the central and temporal macula using Adobe illustrator 2021 (version 25.2.3).

Macula diameters determined by cellular thresholds were made by identifying 5% of the maximum number of cells within the foveal regions of each lizard’s retina. This number was then used to calculate a peripheral retina threshold by adding it to the averages of the outermost 6 bins from the central foveal area (i.e., bins −20, −19, −18, −17, −16 −15 and +15, +16, +17, +18, +19, +20) and the 3 bins from the temporal foveal area (i.e., bins −10, −9, −8 and +8, +9, +10). A smoothing parameter from JMP Pro 14 (*JMP^®^, Version 14 Pro*, 1989–2021) (Smoothing Spline Fit; lambda set at 0.005, standardized) was then applied to the data generated from this calculation. This resulted in a ONL trend line being generated for each of the lizard’s foveal regions. Bins where the ONL trend line was equal to or greater than that individual lizard’s set cellular threshold were identified as a macular bin (Supp. Fig. 3a-b). To account for the absence of photoreceptor cell bodies within the central fovea, bins −5 through +5 were automatically included as macula bins. We then used JMP to add the number of bins together and calculate macula diameter lengths for each lizard.

### 2.8 Graphs, tables, analysis

All graphs were generated using JMP Pro 14 or Prism 9 (*GraphPad Prism, Version 9.4.1*., 1994 - 2022). Tables supporting graphical data were provided showing mean and one standard deviation from the mean for each dataset (i.e., body mass, snout-to-vent length, ocular dimensions, retina length and depth, 20x and 40x cellular counts, and macular diameters). Statistical comparisons between sex or different age groups were performed using nonparametric analyses, Kruskal-Wallis and Mann-Whitney tests (significance level *p*< 0.05). Figures were assembled using a combination of JMP, Prism, Adobe Photoshop and Illustrator. Photoshop was also used to alter white balance and contrast in histological images; in such cases, adjustments were made to the whole image.

## 3. Results and Discussion

### 3.1 Growth Trends

Assessing trends in ocular development first necessitated the assessment of growth patterns of the anole body. Anole lizards grow substantially post-hatching, and it has been noted that male anoles are larger than female anoles. Upon hatching, male and female anoles are similar in size in terms of body mass and snout-to-vent lengths. By adulthood, males are nearly double the weight and have SVLs approximately 20% larger than females (Fig. 2a, c-d). The same was true for their overall eye size (Fig. 2b, e). Statistical analyses (Mann-Whitney tests) showed that there was no significant difference between male and female hatchling body mass (*p* value = 0.7056), snout-to-vent lengths (*p-value* = 0.0881), or eye size (*p* value = 0.2924). By adulthood, however, all three of these metrics exhibited significant differences between males and females (*p-value* < 0.0001 for each metric; Fig. 2c-e). To identify trends in maturity and when male and female anoles diverge, we expanded our initial analyses to incorporate 2-month, 4-month, and 6-month-old juveniles (Supp. Fig 2; Supp. Table 1). By 4 months, female and male body weights were significantly different (*p-value* = 0.0479) (Supp. Fig. 2a). However, differences in SVL (*p-value* = 0.0463) and eye size (*p-value* = 0.0080) were observed as early as 2 months (Supp. Fig. 2b, c). We found that females reached adult weight, SVL, and eye size at close to 6 months (Supp. Fig. 2a-c), around the same period when they also reach sexual maturity and start producing eggs. Males reach sexual maturity closer to 7-8 months in the lab and consistent with this reached adult-size in terms of eye growth, weight, and SVL after 6 months (Supp. Fig. 2a-c).

**Figure 2.**
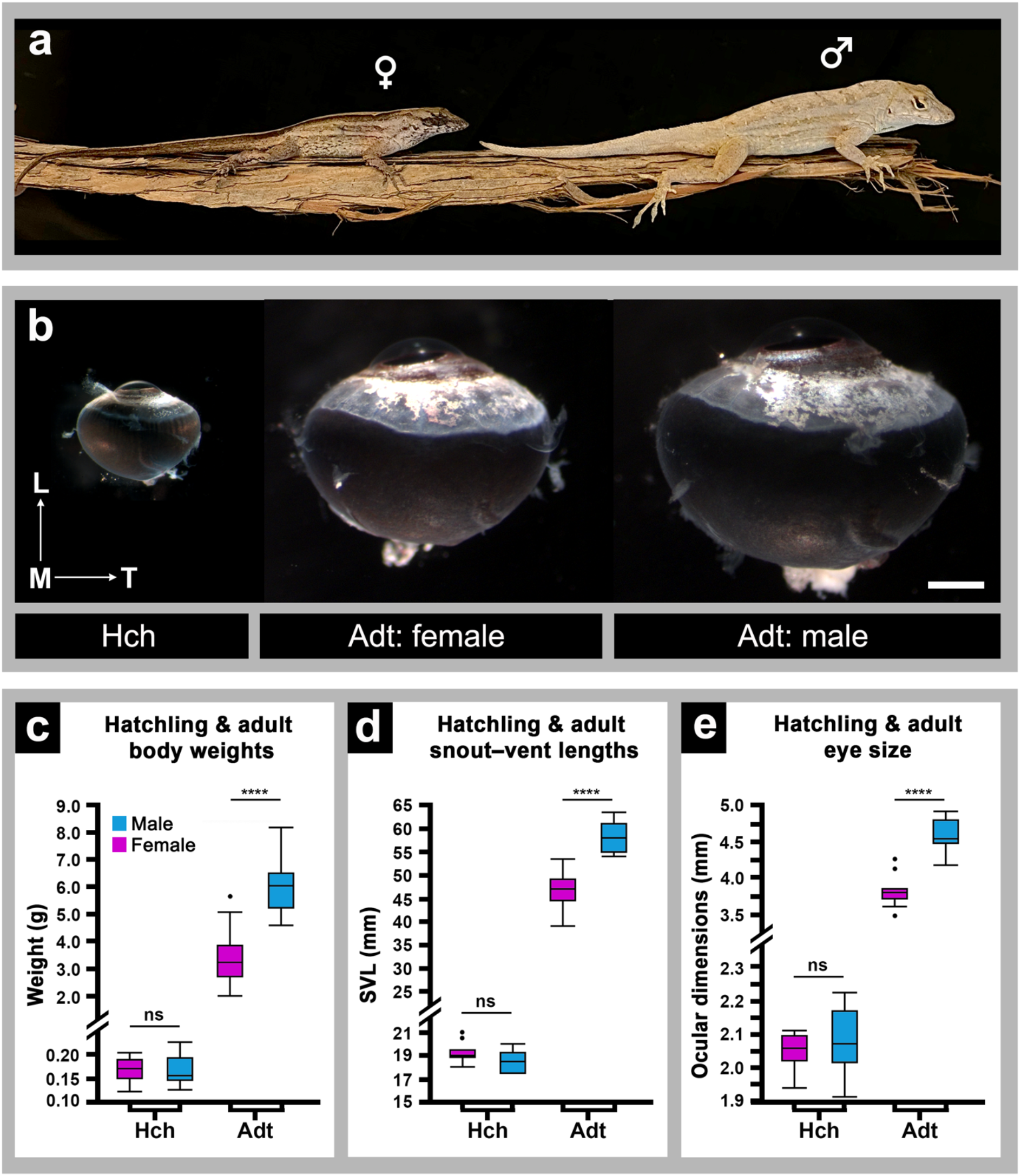
(a) male (right) and female (left) adult lizards, with adult being classified as greater than a year old. (b) Photographs of a hatchling lizard eye (left), an adult female eye (middle), and an adult male eye (right) to scale. The arrow pointing from M to L represents the lateromedial axis while the arrow pointing from M to T represents the temperomedial axis. The scalebar represents 50 microns. (c) Hatchling and adult body weights in grams with sex denoted. The y-axis has a breakpoint that represents the discrepancy between 0.20 grams and 2.0 grams. (d) Hatchling and adult snout to vent lengths measured in millimeters with sex denoted. The y-axis has a breakpoint that represents the discrepancy between 21mm and 25mm. (e) Hatchling and adult ocular dimensions, which are obtained by averaging the three different ocular axes, measured in mm with sex denoted. The y-axis has a breakpoint that represents the discrepancy between 2.3 mm and 3.5 mm.

Ascertaining growth patterns in terms of body size and globe expansion allowed us to assess how retinal measurements were related to the expansion of the globe and body. Further characterization of the retina would allow us to ascertain how pit morphology and photoreceptor cell density changed as a function of retinal and eye growth.

### 3.2 Macular Morphology, Maturation, & Retinal Expansion

We next assessed the morphology of the central and temporal fovea and macula across the age groups (see methods section for additional details regarding definition of macula regions). Specifically, we tracked and quantified changes in foveal shape and size, pit width (i.e., the distance from one parafoveal peak to the other), and pit depth (i.e., the distance from the base of pit to the maximal height of the parafoveal peaks) (See Supp. Table 3 for diagram) and changes in retinal length.

We found that the overall morphology and size of the central fovea remained the same from hatching to adulthood (Fig. 3; Supp Table 2). However, this was not true for the temporal fovea. Instead, it progressively shallows and widens with age and was more variable between sex and individual lizards (Fig. 3). Examining temporal fovea pit dimensions, we found that only pit width, more than depth, was significantly altered, with width doubling in size from hatching to adulthood, with males tending to have slightly broader pits compared to females (Supp. Table 2).

**Figure 3.**
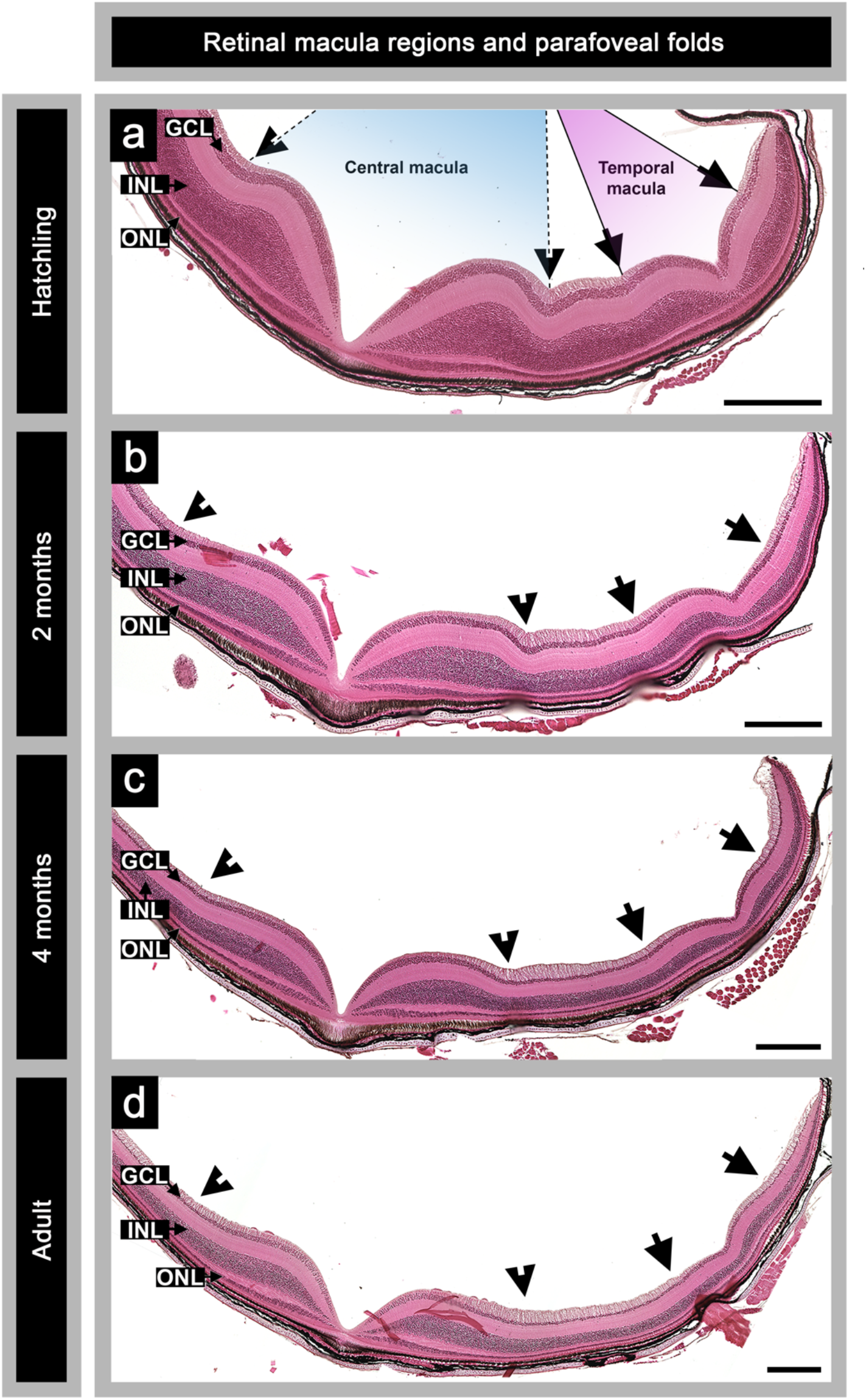
Hematoxylin and eosin stained histological sections from lizards representative of a) hatchlings, b) 2 months, c) 4 months, d) and adults (>1 year). The hatchling panel (A) shows the central macula region between the retinal indentations (arrowheads) highlighted in blue while the temporal macula region between the retinal indentations (arrows) is highlighted in pink. The ganglion cell layer, inner nuclear cell layer, and outer nuclear retinal layers are denoted as GCL, INL, and ONL, respectively. All representative sections are from males and each scale bar represents 200 microns.

We also observed that as the eye grows, the central parafoveal region, its margins, and the shallow depressions which defined the central macula and peripheral retina boundaries become smoother, eventually flattening out (Fig. 3). Similar changes were also observed in the temporal parafoveal region, with the temporal retinal indentations becoming less distinct with age (Fig. 3). Overall, the geometry of both parafoveal regions changed from pronounced indentations to smoother and flatter retinal surfaces. We observed that cells within the macular regions appear to redistribute, potentially driving this change in the retinal landscape (Fig. 3). This finding is particularly pronounced in the inner nuclear layer, and to a lesser extent, in the photoreceptor cell layer (Fig. 3). Photoreceptor cells, which are particularly dense around the central fovea, appear to decrease more rapidly away from the pit centers in the adult compared to hatchlings. In the temporal macular region, all three cell layers appear to decrease in overall thickness or become less dense from hatchling to adulthood (Fig. 3, Fig. 4a-b). These findings, however, are not solely restricted to the temporal macula itself but are evident globally, throughout the peripheral retina areas (Fig. 4b).

**Figure 4.**
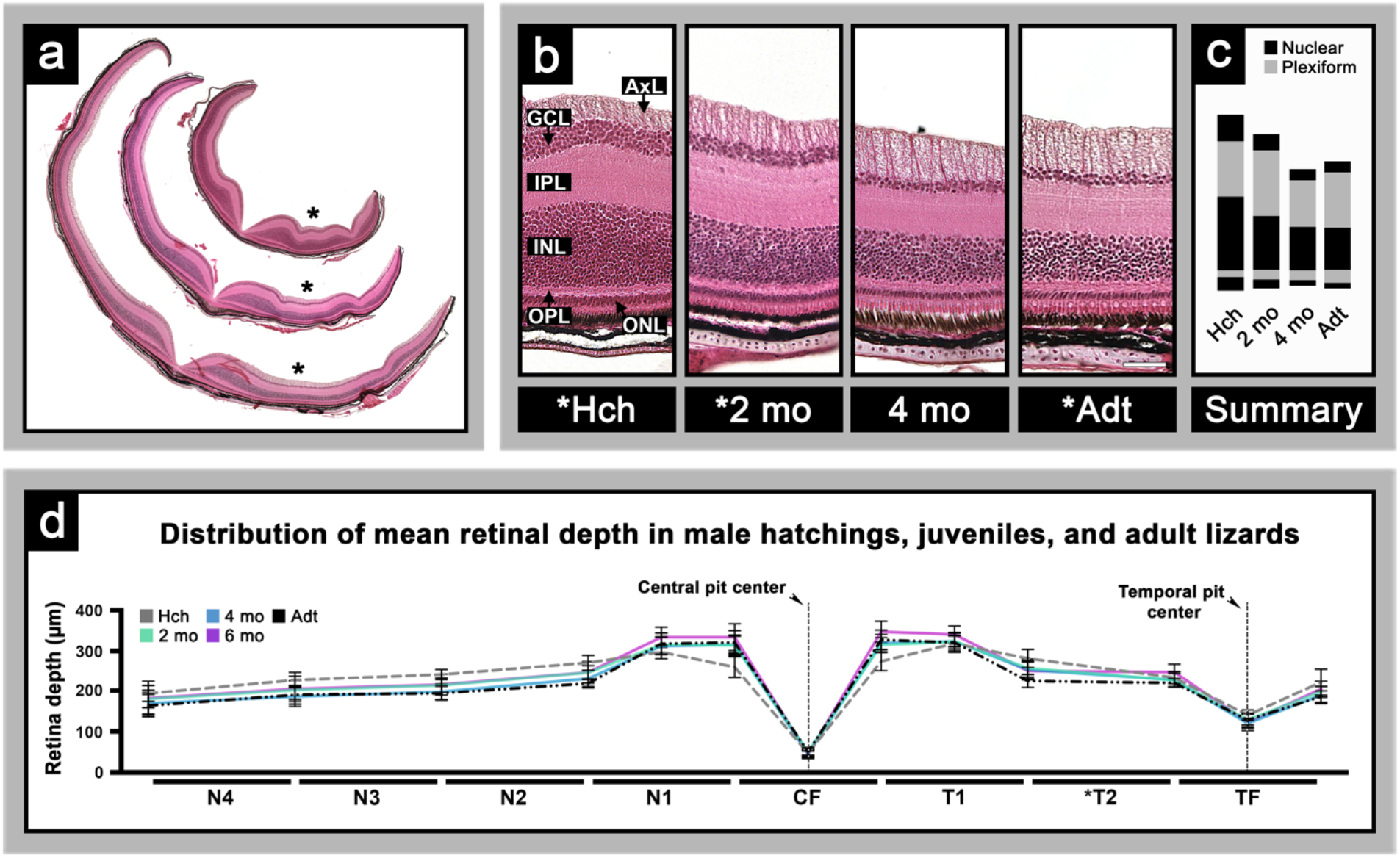
(a) Hematoxylin and eosin-stained histological sections of Hch, 2mo, and Adt lizard eyes scaled to representative sizes in relation to each other. The asterisk represents the location of the T2 bin (b) Magnified T2 bins of the Hch, 2mo, 4mo, and Adt lizard eyes. GCL, IPL, INL, OPL, ONL, and AxL (axon fiber layer) are denoted. (c) Summary of the widths of the nuclear and plexiform layers from the images in Panel B of the Hch, 2mo, 4mo, and Adt lizards. The black rectangles represent the widths of the corresponding nuclear layers while the light gray rectangles represent the widths of the corresponding plexiform layers. (d) Trends in retinal depth across the retinal bins of males from Hch to Adt with central pit and temporal pit centers denoted by the vertical dashed lines. These depths were taken at the lines dividing the retinal bins from one another. These measurements were not taken for the peripheral side of N5 or T4 because the retina tapers together, meaning the retinal width would be 0 micrometers. N1 and T1 occupy a larger region on the x-axis because additional measurements were taken in these bins of the largest retinal depth immediately surrounding the pit. The CF and TF bins also occupy a larger region on the x-axis because additional measurements were taken of the retinal pigmented epithelium to the beginning of the pit to show overall pit depth.

Our data also indicates that ocular growth predominantly occurs through the expansion of the peripheral retina. This is easily seen in the peripheral retina region separating the two foveae, which increases in length and decreases in thickness as ocular growth occurs from hatching to adulthood (Fig. 4a-d, Supp. Table 3). Retinal depth is affected by the thickness of the axonal, nuclear (cellular), and plexiform layers. In the portion of the retina separating the central and temporal foveae (T2 bin), we observed a thinning of the cellular layers concurrently with the overall thinning of the retina (Fig. 4a-c). This suggests to us that as ocular growth occurs, the plexiform and axonal layers also continue to develop (Fig. 4b, c). Overall, we found that most changes occurred in the peripheral retina and areas immediately flanking the central pit, with minimal effect on the foveal regions themselves (Fig. 4d; Supp. Table. 3).

Our observations here are consistent with investigations in primates that showed that the peripheral retina thins to a greater degree than central retina (Springer & Hendrickson, 2004b) as well as other studies in vertebrates where retina thinning was seen after peripheral retina expansion (Kuhrt et al., 2012; Mastronarde et al., 1984; Reichenbach et al., 1991). Optical coherence tomography scans in birds have also shown that pit depth tended to be deeper in juveniles compared to older individuals (Potier et al., 2020). These findings, along with the alterations to foveae morphology and the loss of the retinal indentations, occur concurrently with eye growth in terms of retinal length and ocular dimensions. This offers evidence for growth induced retinal remodeling (Springer & Hendrickson, 2004a, 2004b, 2005). Changes in cell distribution would also seem to support this. Interestingly, however, the decrease in photoreceptor cell density immediately adjacent to the foveal pits is contrary to what has been observed in the fovea of primates, where photoreceptor cells increase in number as the eye grows (Hendrickson et al., 2012b; Springer & Hendrickson, 2005; Yuodelis & Hendrickson, 1986).

### 3.3 Retina Cell Density

We next wanted to assess how eye size might influence cell distribution in the retina of the aging anole. If retinal length increases over time, and cell number remains constant, this would provide evidence for retinal stretch influencing retinal remodeling and macular maturation. To investigate this, we generated a pipeline that performed automated cell counts across retinal sections (Supp. Fig. 1b; see methods section for additional details) and compared the total cell numbers to eye size and retina lengths. We found that anoles at the time of hatching had the highest cell density (GCL + INL + ONL) per plane of section for all age groups (Fig. 5a; Supp. Table 4). Over time, these numbers decreased, reaching the lowest observed value by adulthood for both sexes (Fig. 5a, c).

**Figure 5.**
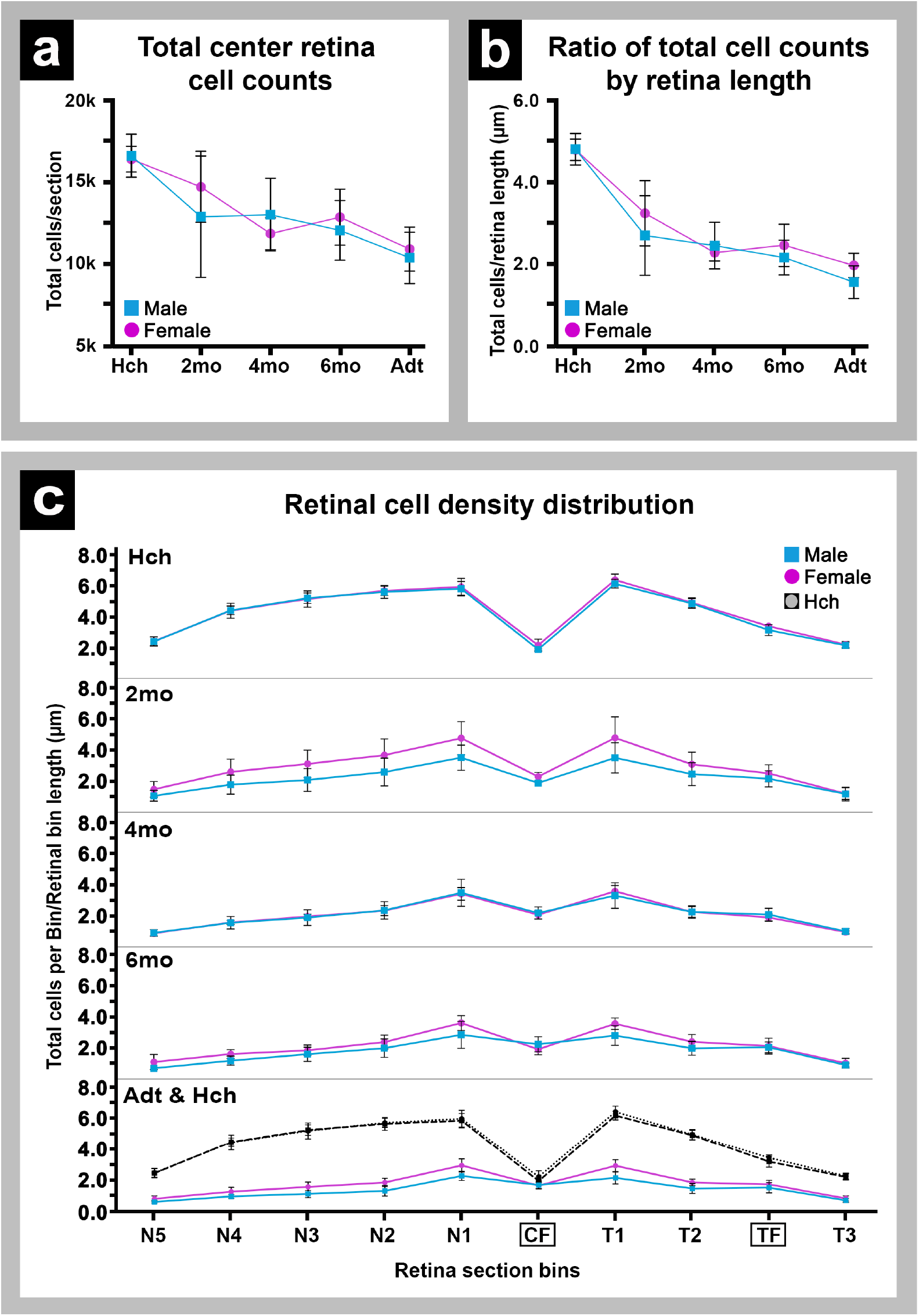
(a) Total cell counts (GCL+INL+ONL) on representative retinal sections of male and female lizards across age groups (Hch, 2mo, 4mo, 6mo, Adt). (b) Cell densities (cells/μm) of retinal sections measured by taking the total cell counts (GCL+INL+ONL) divided by retinal length of male and female lizards across age groups (Hch, 2mo, 4mo, 6mo, Adt). Panels a and b have female lizards denoted by pink boxplots and male lizards denoted by blue boxplots. (c) Cell density across the retina, assessed by taking total cell counts per bin divided by retinal length per bin for each sex and age group. Females are represented by the pink distribution and individual datapoints are represented by the circles. Males are represented by the blue distribution and individual datapoints are represented by the triangles. Hch data points, represented by the black dashed lines, are overlaid on the Adt graph for comparison (bottom panel).

A drop in cellularity from hatchling to adulthood raises the question of whether cell death is occurring within the retina. Studies in embryonic rodents, chicks, turtles and anoles have shown that a significant portion of cell death occurs in retina cell layers during development, but that cell death slows and largely ceases shortly after birth or hatching (Beazley et al., 1987; Cook et al., 1998; Francisco-Morcillo et al., 2004; Rasys, Pau, Irwin, Luo, Kim, Wahle, Trainor, et al., 2021; Rasys, Pau, Irwin, Luo, Kim, Wahle, Menke, et al., 2021). Although we did not specifically look for markers of cell death in retinal sections, we did not observe any evidence of pyknotic cells within our sections across the age groups, suggesting that if cell death is occurring, it is likely minimal. Rather, this decrease in cell numbers is likely an artifact of our plane of section. For instance, if we assume that anoles at the time of hatching possess a full complement of cells and that number remains largely constant, then as the eye gets larger, there will be a decreased density of cells across each section due to the expansion of the retinal tissue. Hence, a more compact eye would lead to higher cell numbers, while a larger eye with the same number of cells would have a reduced number of cells per section. Our results also indicate that ocular growth and retinal remodeling are not influenced by an increase in retinal cell number or size, which is not surprising as investigations in a variety of organisms have found that retinal cells have low proliferative activity in the maturing retina (Alvarez-Hernan et al., 2021; Diaz-Araya & Provis, 1992; Rapaport et al., 2004). While it is possible that there is some residual cell division activity in the anole retina, our data indicate that cell redistribution likely plays a much larger role in retinal remodeling and growth and can account for the decline in cell numbers from hatchlings to adults. Together these findings lend strength to the model that growth-induced retinal stretch, as suggested in primates (Springer & Hendrickson, 2005), may be an underlying factor in retina and macula remodeling.

### 3.4 Macular Photoreceptor Cell Distribution & Diameters

We next wanted to look more closely at photoreceptor cell distribution within the macula and the diameter of the macula itself to determine if it changes over time. Historically, in humans as well as other primates, changes to the macula have been assessed by measuring the rod-free zone (Yuodelis & Hendrickson, 1986). However, anoles possess a pure coned retina, lacking a rod-free zone (Fite & Lister, 1981; Walls, 1942), making it impossible to use this as a metric to define boundaries. Moreover, the anole retina is avascular (Fite & Lister, 1981; Rasys, Pau, Irwin, Luo, Kim, Wahle, Trainor, et al., 2021), also precluding the use of the avascular zone present in primates to assess modifications to the macula (Eldaly et al., 2020; Hussain & Hussain, 2016). Therefore, to examine subtle changes to photoreceptor cell distribution more thoroughly in anoles, we performed cell counts on sections imaged at a higher magnification (40x) using the criteria outlined in Methods. This time, however, foveal areas were binned and surveyed by calculating the macula distance in optical degrees, normalizing to each lizard’s individual axial measurements (Supp. Fig. 1b; see methods section for additional details). Doing so allowed us to account for differences in eye size as well as accurately tailor the distance of retina required to properly assess the macula regions of both the central and temporal foveae. Because total retina cell counts were not dramatically different for males and females, data from both sexes were evaluated together for each age group.

Our analysis revealed that the highest concentrations of photoreceptor cells were located around the central and temporal foveae. Cell density gradually decreased as one moved away from either pit center (Fig. 6a-b). When we looked for differences in cell distribution between hatchlings and adults, we found that the peak number of photoreceptor cells in the central parafoveal area increased from 30 to 50 cells/bin (Fig 6a). Similarly, cell numbers in the temporal fovea changed from 20 to 28 cells/bin, whereas at the peripheral-most boundaries, cell numbers largely stayed the same, ranging between 10 to 20 cells/bin (Fig 6a-b). When we calculated cellular density, we found that peak cell numbers around the central fovea were actually relatively constant, approximately 0.8 cells/μm across all age groups. However, the locations of peak cell numbers shifted over time. In hatchlings, the cell peaks around the central fovea are located between −7 and +6 degrees away from the central fovea but were within 4 degrees from the pit center in adults (Fig 6c; Supp. Table 5). We also observed that the location of the retinal region with inner nuclear layer peak cell numbers moved closer to the central fovea pit center with age (Supp. Fig. 4; Supp. Table 6). This was not apparent in the ganglion cell layer or observed in any of the retinal cell layers within the temporal macula region (Supp. Fig. 4; Supp. Table 7). The most startling difference, however, was the observation that both the temporal macular cell number peak and peripheral cell density dropped (Fig. 6d; Supp. Table 5). In hatchlings, temporal peak cell density was around 0.5 cells/μm, while in adults, this value decreased to 0.3-0.4 cells/μm; a similar degree of change was observed at peripheral margins for both macular areas (Fig 6d). Taken together, this data suggests that some level of cell packing occurs within the anole retina post-hatching.

**Figure 6.**
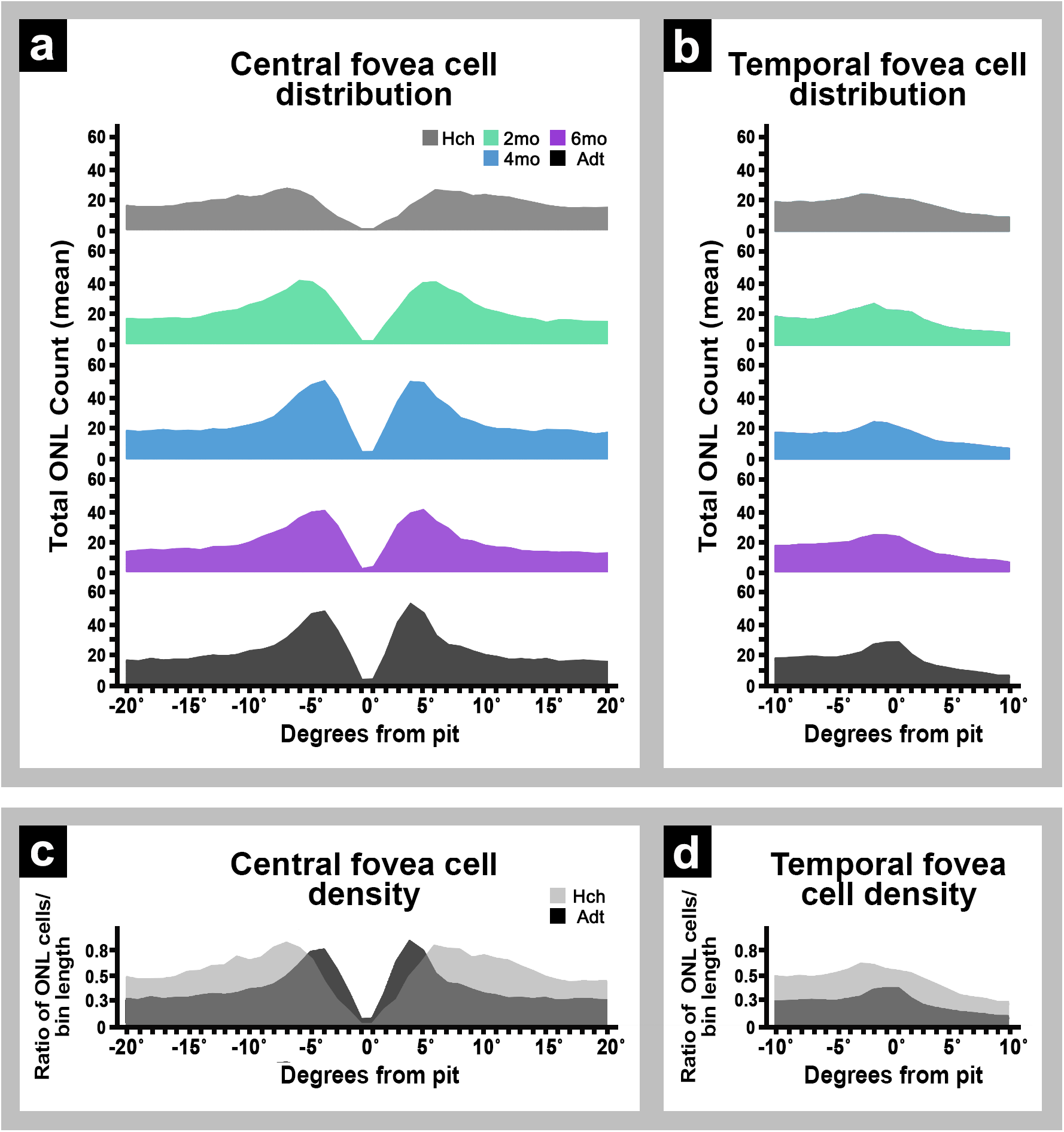
(a) Central fovea cell distribution as measured by mean ONL cell counts within 20 degrees on either side of the center foveal pit. The negative degrees represent the retina extending on the nasal side of the center foveal pit while the positive degrees indicate the retina extending on the temporal side of the center foveal pit, closer to the temporal fovea. (b) Temporal fovea cell distribution as measured by mean ONL cell counts within the 10 degrees on either side of the temporal foveal pit. The negative degrees represent the intrafoveal region, which is the retina between the central and temporal fovea pits. The positive degrees indicate the marginal retina, which is the retina on the other side of the temporal fovea pit that is closer to the ciliary marginal zone and cornea. (c) Central fovea cell density as measured by the number of cells in the ONL per bin divided by the respective bin length extending 20 degrees on either side of the center foveal pit. The light gray color represents the hatchling distribution while the dark gray color represents the adult distribution. (d) Temporal fovea cell density as measured by the number of cells in the ONL per bin divided by the respective bin length extending 10 degrees on either side of the temporal foveal pit. The light gray color represents the hatchling distribution while the dark gray color represents the adult distribution.

When we looked for changes in macular diameter using either visual assessments or a non-biased cellular threshold-based approach (Supp. Fig. 3a; see methods for more details), we found that both central and temporal maculae increased in diameter between hatchlings and adults, and, although subtle, macular expansion in males was a slightly greater than in females (Supp. Fig. 3c; Supp. Table 2). However, when we compared eye size, each macula comprised a smaller fraction of the total retinal landscape as eye size increased, and no difference in sex was observed (Supp. Fig. 3d). We also found that both central and temporal maculae in females reach near adult dimensions at around 4 months, while males reached this value sometime after 6 months (Supp. Fig. 3c), matching roughly with the timeline of when females and males mature in terms of SVL, ocular averages, weight, and sexual maturity. Studies in primates have shown that the diameter of the rod-free zone in the retina decreases, suggesting a decrease in the macular diameter (Yuodelis & Hendrickson, 1986). Although not directly applicable because the anole lacks a rod-free zone, our results suggest that macular dimensions do not change as a function of ocular growth in anoles.

Similarly, our data suggest that inner nuclear layer and photoreceptor cell packing during periods of ocular growth after hatching in anoles is rather minimal and concludes roughly around 2 months (Supp. Fig. 4), contrary to what has been observed in primates (Springer & Hendrickson, 2005; Springer et al., 2011; Yuodelis & Hendrickson, 1986). Instead, foveal areas seem to undergo more of a refinement process. Cells immediately adjacent to the central pit are preserved – or in the case of the temporal fovea, become slightly reduced in number – whereas cells located near the outer central and temporal parafoveal margins gradually decrease in number over time (Fig. 6c-d). This decrease in cell density in the temporal retina is consistent with the finding that retinal thickness also decreases most substantially in the peripheral retinal regions. It is possible that because the temporal fovea is located near the ciliary marginal zone, that these regions of the eye expand at a faster rate than the central retina and, subsequently, may undergo a more extensive remodeling process. Our axial measurements of the anole eye would support this (Supp. Table 1). Another possible explanation is that during embryonic development anoles experience dynamic changes in ocular shape (Rasys, Pau, Irwin, Luo, Kim, Wahle, Trainor, et al., 2021). Foveal areas undergo periods of localized elongation followed by retraction. During ocular retraction, pit formation and the majority of photoreceptor packing occurs (Rasys, Pau, Irwin, Luo, Kim, Wahle, Trainor, et al., 2021; Rasys, Pau, Irwin, Luo, Kim, Wahle, Menke, et al., 2021). It is possible that ocular retraction, particularly along the z-axis, may persist for a brief period after hatching and during ocular expansion. This could result in the generation of inward forces within the central macular region and thus provide a mechanism for the refinement that we see in the cell populations around the central fovea. Regardless, these findings argue that cell redistribution is likely the major underlying factor accommodating retinal expansion as eye size increases.

### 3.5 Concluding remarks

In primates, it has been proposed that cone photoreceptor cells pack more densely into the center of the macula and that, as a result, the diameter of the rod-free zone within the macula decreases (Patel et al., 2017; Provis & Hendrickson, 2008; Yuodelis & Hendrickson, 1986). Springer and Hendrickson’s hypothesis for macula and fovea maturation in primates is that extrinsic forces (i.e., retinal stretch, intraocular pressure) generated from ocular growth act on cells, and the cells passively respond by packing more tightly into the center of the macula, thus decreasing the diameter of the rod free zone within the macula (Springer & Hendrickson, 2005). Our data does not appear to support this hypothesis in anoles. Macular dimensions actually increase in size and although photoreceptor cell and inner nuclear layer packing (around the central fovea) does occur between hatching to 2 months of age, this is almost negligible and does not continue throughout the remaining periods of ocular growth in anoles. Rather, our work suggests that at the time of hatching lizards already have a full complement of cells within each macula and foveal maturation occurs predominantly through the redistribution of its outermost cells to accommodate peripheral retina expansion.

This work provides a comprehensive analysis of retinal maturation and sex differences in anoles. Now that genetic tools are available, anoles are an exciting model to test how retinal maturation patterns are established (Rasys et al., 2019).

## Code availability

The code used for cell quantification is available: https://github.com/hannahqkim/CellCountProtocol

## Acknowledgements

M.A.W. and H.Q.K contributed equally to the study conception, data collection, analysis, interpretation of results, and draft manuscript preparation. A.M.R. contributed to study conception, design, analysis, interpretation of results and draft manuscript preparation, while D.B.M and J.D.L. contributed to contributed to study conception and draft manuscript preparation. This work was funded through NSF #1827647 awarded to D.B.M and J.D.L as well as through the Center for Undergraduate Research Opportunities (CURO) at the University of Georgia.

