## Supplemental Material for "Maturation and Refinement of the Maculae and Foveae in the Anolis sagrei lizard"

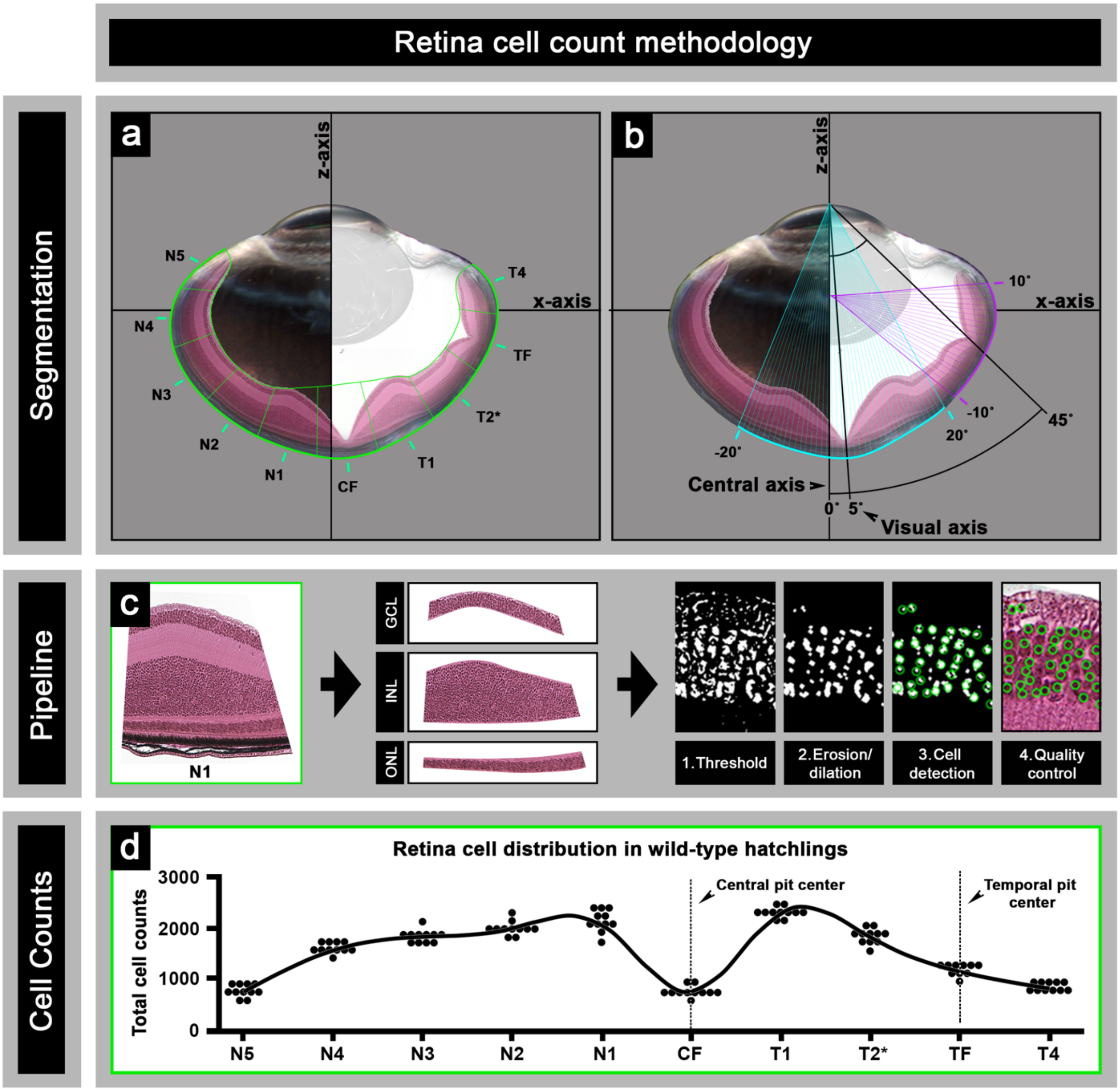


#### Supplemental Figure 1

(a) Methodology for performing 20x cell counts using (retinal length/10) to split the retina into 10 bins. To center the CF and TF bins on center foveal pit and temporal foveal pit, 5 nasal bins (N5-N1) and 4 temporal bins (T1-T2, TF, T4) were designated. (b) Methodology for performing 40x cell counts using optical degrees with the origins of each macula at pit center. The central fovea is located approximately 5˚ (denoted as the visual axis) temporally away from central axis (defined as 0°), whereas the center of temporal pit is roughly positioned 45° temporal of this central axis. The z-axis was used to calculate retinal arc lengths corresponding to 20 degrees on either side of the pit for the central macula. The x-axis was used to calculate retinal arc lengths corresponding to 10 degrees on either side of the pit for the temporal macula. (c) An overview of the steps in the pipeline for performing cell counts, including separating the retinal layers, thresholding, erosion / dilation, cell detection, and applying a quality control. (d) Total cell counts (GCL+INL+ONL) in the 10 20x bins in Hch lizards, with central pit and temporal pit center denoted by a dashed line.

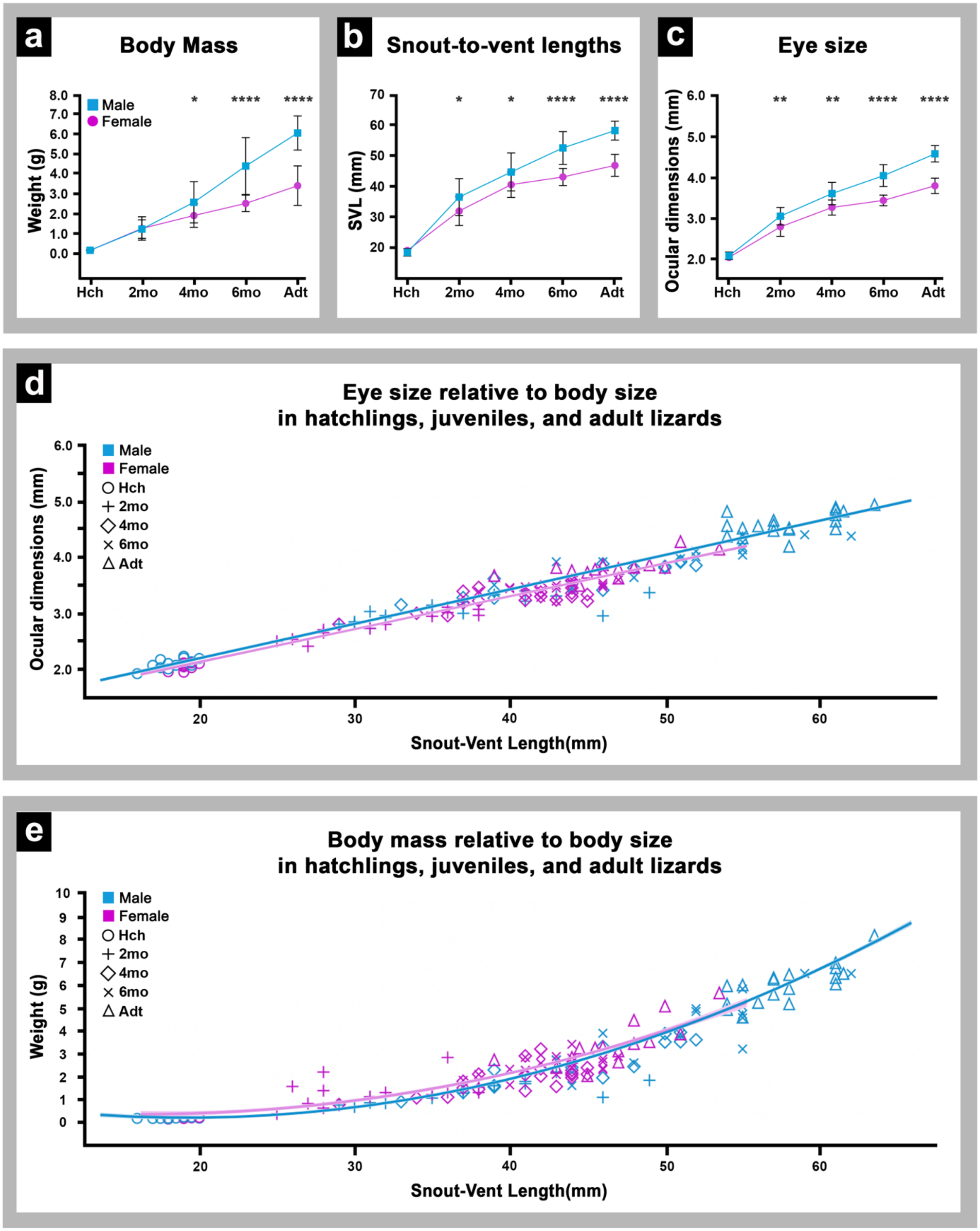


**Supplemental Figure 2**

(a) Body weights in grams of males and females in comparison to age (Hch, 2mo, 4mo, 6mo, Adt). (b) Snout to vent lengths in mm of males and females in comparison to age (Hch, 2mo, 4mo, 6mo, Adt).

(c) Ocular dimensions in mm of males and females in comparison to age (Hch, 2mo, 4mo, 6mo, Adt). Panels A-C show female data in pink boxplots and male data in blue boxplots. (d) Ocular dimensions in millimeters compared to snout to vent length in millimeters across the age groups with sex denoted by color and age group denoted by shape. The equation for the line of best fit that is merged between the sexes is $y = 59.82x + 934.6$, which can be used to calculate predicted ocular dimensions and to form a rough estimate of juvenile age. (e) Body weight in grams compared to snout to vent length in millimeters across the age groups with sex denoted by color and age group denoted by shape. The equation for the line of best fit that is merged between the sexes is $y = 1.29 - .1219x + 0.00344x^2$, which can be used to calculate predicted weight and to form a rough estimate of juvenile age.

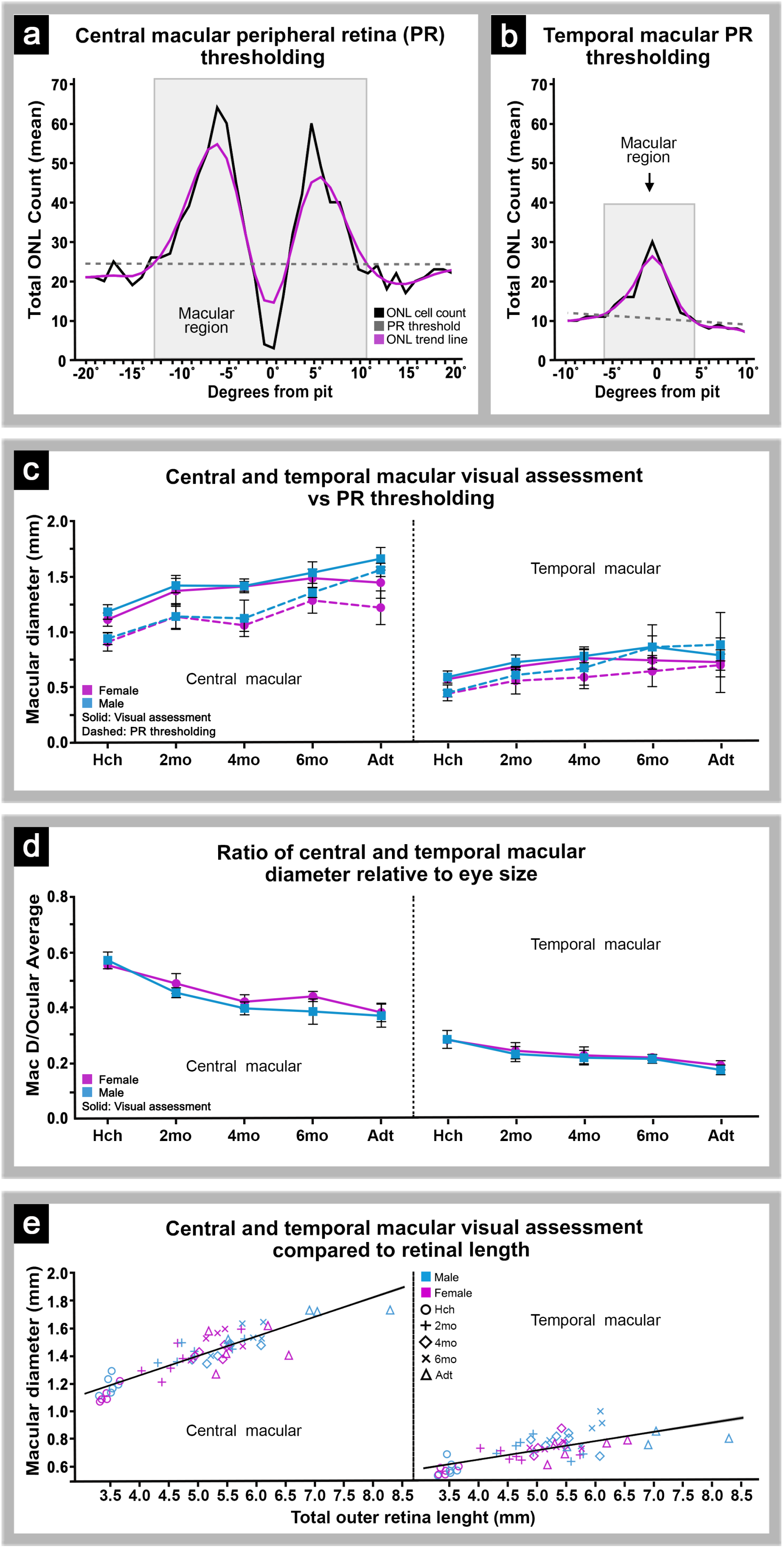

a
Central macular peripheral retina (PR) thresholding
Total ONL Count (mean)
Macular region
ONL cell count
PR threshold
ONL trend line
Degrees from pit
b
Temporal macular PR thresholding
Macular region
Degrees from pit
c
Central and temporal macular visual assessment vs PR thresholding
Macular diameter (mm)
Temporal macular
Central macular
Female
Male
Solid: Visual assessment
Dashed: PR thresholding
Hch
2mo
4mo
6mo
Adt
d
Ratio of central and temporal macular diameter relative to eye size
Mac D/Ocular Average
Temporal macular
Central macular
Female
Male
Solid: Visual assessment
e
Central and temporal macular visual assessment compared to retinal length
Macular diameter (mm)
Male
Female
Hch
2mo
4mo
6mo
Adt
Temporal macular
Central macular
Total outer retina lenght (mm)

**Supplemental Figure 3**

(a) An example of the cell-count based 5% threshold based on peripheral retina applied to the central macula. The black line represents the ONL cell count distribution within the degrees surrounding the pit, the pink line represents the smoothing curve applied by JMP, and the gray region represents bins identified as being within the macula based on the dashed horizontal line representing the 5% threshold. (b) An example of the cell-count based 5% threshold based on peripheral retina applied to the temporal macula. The black line represents the ONL cell count distribution within the degrees surrounding the pit, the pink line represents the smoothing curve applied by JMP, and the gray region represents bins identified as being within the macula based on the dashed horizontal line representing the 5% threshold. (c) Central and temporal macular diameters as determined by both visual assessment and cell-count thresholding across the age groups. (d) The ratio of central and temporal macular diameter relative to eye size as determined by taking macular diameter divided by ocular averages across the sexes and age groups. (e) Central and temporal macular diameters (mm) plotted against total outer retinal length (mm) with age and sex denoted.

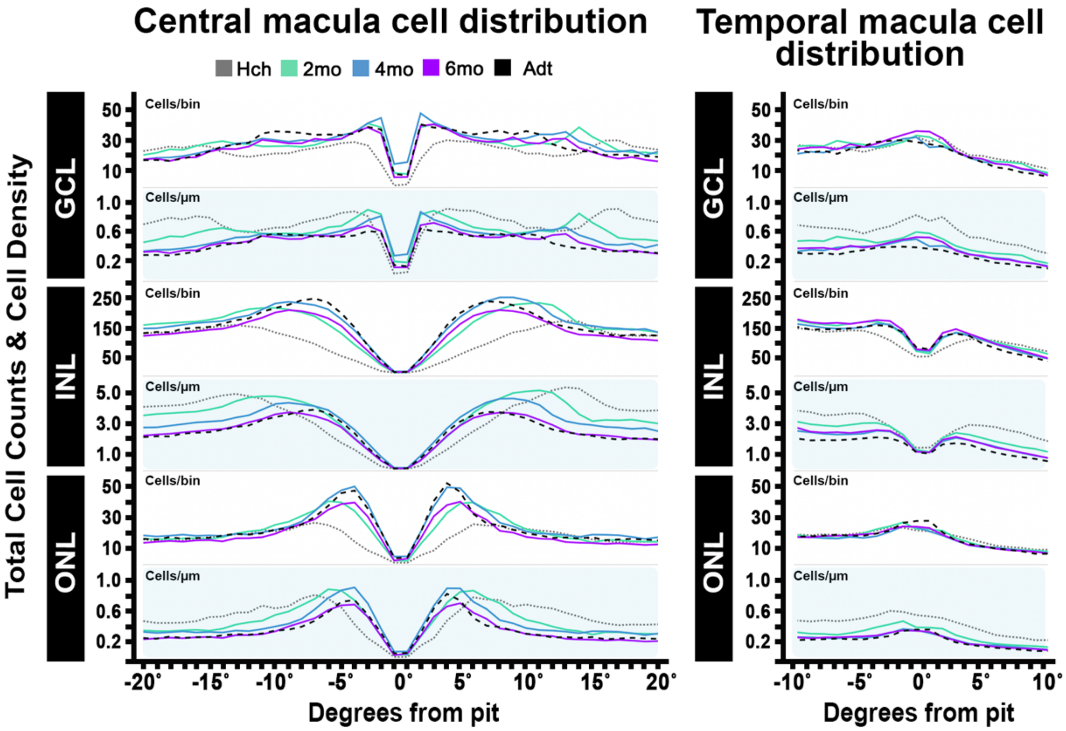


**Supplemental Figure 4**

Central and temporal macula cell distribution for all three cellular layers of the retina – the GCL, INL, and ONL— with age (Hch, 2mo, 4mo, 6mo, Adt) denoted by color of the line. Both total cells/bin and cell density (cells//µm) are shown graphically. For the central fovea, the negative degrees represent the optical degrees on the retina extending on the nasal side of the center foveal pit while the positive degrees indicate the optical degrees on the retina extending on the temporal side of the center foveal pit, closer to the temporal fovea. For the temporal fovea, the negative degrees represent the intrafoveal region, which is the retina between the central and temporal fovea pits. The positive degrees indicate the marginal retina, which is the retina on the other side of the temporal fovea pit that is closer to the ciliary marginal zone and cornea.

| SEX | AGE | n | Weight (g)<br>Mean ± SD | SVL (mm)<br>Mean ± SD | Ocular Dimensions (µm)<br>Mean ± SD | X-axis (µm)<br>Mean ± SD | Y-axis (µm)<br>Mean ± SD | Z-axis (µm)<br>Mean ± SD |
|---|---|---|---|---|---|---|---|---|
| Female | Hch | 25 | 0.17 ± 0.03 | 19.1 ± 0.7 | 2051 ± 54 | 2188 ± 77 | 2034 ± 68 | 1817 ± 53 |
| | 2 mo | 15 | 1.32 ± 0.7 | 31.93 ± 4.6 | 2808 ± 231 | 3084 ± 217 | 2909 ± 231 | 2466 ± 184 |
| | 4 mo | 21 | 1.91 ± 0.59 | 40.52 ± 4.2 | 3271 ± 186 | 3661 ± 139 | 3633 ± 160 | 2881 ± 130 |
| | 6 mo | 21 | 2.52 ± 0.41 | 43 ± 2.8 | 3445 ± 132 | 657 ± 152 | 3457 ± 160 | 3014 ± 112 |
| | Adt | 15 | 3.41 ± 0.99 | 46.7 ± 3.5 | 3803 ± 188 | 4080 ± 179 | 3880 ± 219 | 3380 ± 163 |
| Male | Hch | 20 | 0.16 ± 0.03 | 18.35 ± 1.0 | 2081 ± 91 | 2238 ± 57 | 2089 ± 70 | 1886 ± 56 |
| | 2 mo | 17 | 1.23 ± 0.46 | 36.47 ± 6.0 | 3059 ± 213 | 3414 ± 244 | 3262 ± 294 | 2737 ± 149 |
| | 4 mo | 14 | 2.57 ± 1.04 | 44.64 ± 6.2 | 3614 ± 274 | 3825 ± 331 | 3629 ± 279 | 3292 ± 267 |
| | 6 mo | 15 | 4.39 ± 1.42 | 52.47 ± 5.3 | 4053 ± 268 | 4352 ± 294 | 4118 ± 219 | 3556 ± 244 |
| | Adt | 19 | 6.04 ± 0.86 | 58.05 ± 3.0 | 4589 ± 200 | 4910 ± 262 | 4601 ± 238 | 3998 ± 201 |

**Supplemental Table 1**

Mean weight (grams), snout to vent length (mm), ocular dimensions (µm), and x-. y-, and z—axes (µm) across sexes and age groups.

| SEX | AGE | n | Retina length (μm) | CF Macular Diameter (Mean ± SD in μm) Visual Assessment of Macular Diameter | Macular Diameter by 5% Threshold | Pit Depth | Pit Width | TF Macular Diameter (Mean ± SD in μm) Visual Assessment of Macular Diameter | Macular Diameter by 5% Threshold | Pit Depth | Pit Width |
|---|---|---|---|---|---|---|---|---|---|---|---|
| Femal | Hch | 5 | 3454 ± 134 | 1116 ± 60 | 913 ± 83 | 325 ± 13 | 551 ± 17 | 573 ± 27 | 443 ± 31 | 107 ± 11 | 240 ± 9 |
| | 2mo | 6 | 4668 ± 568 | 1376 ± 140 | 1143 ± 107 | 324 ± 22 | 564 ± 83 | 686 ± 33 | 560 ± 120 | 120 ± 15 | 373 ± 72 |
| | 4mo | 4 | 5216 ± 265 | 1417 ± 45 | 1065 ± 55 | 325 ± 17 | 605 ± 69 | 764 ± 83 | 592 ± 72 | 121 ± 22 | 434 ± 18 |
| | 6mo | 6 | 5367 ± 312 | 1491 ± 82 | 1289 ± 114 | 351 ± 41 | 642 ± 40 | 744 ± 19 | 646 ± 141 | 158 ± 25 | 429 ± 40 |
| | Adt | 5 | 5664 ± 559 | 1451 ± 141 | 1224 ± 153 | 309 ± 23 | 578 ± 31 | 727 ± 73 | 700 ± 247 | 134 ± 13 | 466 ± 69 |
| Male | Hch | 6 | 3508 ± 110 | 1185 ± 65 | 944 ± 55 | 330 ± 12 | 573 ± 50 | 592 ± 55 | 449 ± 72 | 117 ± 8 | 237 ± 12 |
| | 2mo | 7 | 4994 ± 535 | 1425 ± 65 | 1144 ± 117 | 326 ± 25 | 605 ± 47 | 728 ± 65 | 611 ± 77 | 112 ± 21 | 400 ± 43 |
| | 4mo | 6 | 5399 ± 393 | 1421 ± 62 | 1127 ± 165 | 325 ± 17 | 582 ± 63 | 782 ± 59 | 677 ± 189 | 133 ± 33 | 446 ± 35 |
| | 6mo | 5 | 5803 ± 349 | 1542 ± 98 | 1361 ± 45 | 330 ± 31 | 583 ± 75 | 866 ± 99 | 865 ± 201 | 138 ± 58 | 468 ± 33 |
| | Adt | 4 | 6905 ± 920 | 1667 ± 103 | 1565 ± 60 | 321 ± 21 | 620 ± 71 | 790 ± 50 | 1027 ± 94 | 109 ± 38 | 555 ± 63 |

**Supplemental Table 2**

Central and temporal macular diameters reported both by visual assessment and by applying a cell-count based 5% threshold. Macular diameters are reported by age group and by sex with overall retinal length noted. The central and temporal foveal pit depth and width as determined by visual assessment are also reported by age group and sex.

### Retinal measurements

#### Outer Retinal Length (Mean ± SD in μm)

| SEX | AGE | n | N5 | N4 | N3 | N2 | N1 | CF | T1 | T2 | TF | T4 |
|---|---|---|---|---|---|---|---|---|---|---|---|---|
| Female | Hch | 5 | 295 ± 42.1 | 345 ± 13.4 | 345 ± 13.4 | 345 ± 13.4 | 345 ± 13.4 | 345 ± 13.4 | 362 ± 17.8 | 362 ± 17.2 | 343 ± 13.9 | 358 ± 21.5 |
| | 2 mo | 6 | 477 ± 125.3 | 467 ± 56.8 | 467 ± 56.8 | 467 ± 56.8 | 467 ± 56.8 | 467 ± 56.8 | 463 ± 59.4 | 451 ± 55.7 | 467 ± 58.5 | 482 ± 62.4 |
| | 4 mo | 4 | 612 ± 45.6 | 522 ± 26.5 | 522 ± 26.5 | 522 ± 26.5 | 522 ± 26.5 | 522 ± 26.5 | 503 ± 55.6 | 499 ± 51 | 521 ± 26 | 479 ± 38.7 |
| | 6 mo | 6 | 558 ± 108.6 | 537 ± 31.2 | 537 ± 31.2 | 537 ± 31.2 | 537 ± 31.2 | 537 ± 31.2 | 535 ± 30.8 | 529 ± 32.3 | 529 ± 37.3 | 488 ± 55.5 |
| | Adt | 6 | 702 ± 70.1 | 566 ± 55.9 | 566 ± 55.9 | 566 ± 55.9 | 566 ± 55.9 | 566 ± 55.9 | 559 ± 50.5 | 544 ± 30.7 | 561 ± 63.8 | 522 ± 123.2 |
| Male | Hch | 6 | 294 ± 34.7 | 351 ± 11 | 351 ± 11 | 351 ± 11 | 351 ± 11 | 351 ± 11 | 369 ± 17.8 | 369 ± 19.9 | 346 ± 12.1 | 354 ± 35 |
| | 2 mo | 7 | 546 ± 117.7 | 499 ± 53.5 | 499 ± 53.5 | 499 ± 53.5 | 499 ± 53.5 | 499 ± 53.5 | 483 ± 44.8 | 476 ± 35.2 | 491 ± 46.4 | 519 ± 65.8 |
| | 4 mo | 6 | 623 ± 116.4 | 540 ± 39.3 | 540 ± 39.3 | 540 ± 39.3 | 540 ± 39.3 | 540 ± 39.3 | 529 ± 26.2 | 530 ± 25.7 | 541 ± 40.2 | 530 ± 55.2 |
| | 6 mo | 5 | 693 ± 112.7 | 580 ± 34.9 | 580 ± 34.9 | 580 ± 34.9 | 580 ± 34.9 | 580 ± 34.9 | 588 ± 42.2 | 573 ± 46.6 | 565 ± 37.9 | 578 ± 56.1 |
| | Adt | 5 | 847 ± 89.4 | 665 ± 63.1 | 665 ± 63.1 | 665 ± 63.1 | 665 ± 63.1 | 665 ± 63.1 | 660 ± 60 | 642 ± 51.3 | 654 ± 60.4 | 613 ± 59.3 |

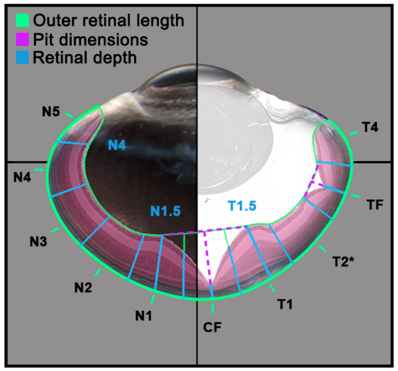


#### Retinal Depth (Mean ± SD in μm)

| SEX | AGE | n | N5 | N4 | N3 | N2 | N1 | N1.5 | CF | Pit Center | T1 | T1.5 | T2 | TF | Pit Center | T4 |
|---|---|---|---|---|---|---|---|---|---|---|---|---|---|---|---|---|
| Female | Hch | 5 | - | 180 ± 9.4 | 213 ± 8.7 | 231 ± 8.5 | 260 ± 12.9 | 291 ± 20.8 | 252 ± 32.6 | 45 ± 4.5 | 269 ± 29 | 311 ± 23.8 | 268 ± 17.6 | 229 ± 13.1 | 137 ± 10.2 | 203 ± 16.6 |
| | 2 mo | 6 | - | 189 ± 22.1 | 204 ± 28.6 | 213 ± 25 | 245 ± 21.4 | 309 ± 16.3 | 312 ± 31.2 | 53 ± 8.8 | 310 ± 16.9 | 320 ± 14.9 | 260 ± 28.7 | 228 ± 22.8 | 126 ± 15.8 | 199 ± 18.9 |
| | 4 mo | 4 | - | 180 ± 20.7 | 193 ± 23.9 | 205 ± 12.1 | 232 ± 11.9 | 310 ± 28.9 | 313 ± 27.1 | 51 ± 8.7 | 312 ± 21.3 | 318 ± 13.1 | 250 ± 15.3 | 219 ± 17.7 | 123 ± 15.5 | 181 ± 19.5 |
| | 6 mo | 6 | - | 196 ± 53.6 | 216 ± 35.5 | 217 ± 30.4 | 247 ± 29.3 | 329 ± 23.8 | 320 ± 29.2 | 45 ± 10.3 | 341 ± 26.2 | 337 ± 15.4 | 256 ± 17.8 | 246 ± 25.1 | 116 ± 24.3 | 201 ± 22.4 |
| | Adt | 6 | - | 169 ± 37.5 | 190 ± 30.7 | 196 ± 22.9 | 225 ± 11.6 | 306 ± 25.2 | 303 ± 27.1 | 49 ± 13.1 | 315 ± 21.8 | 312 ± 27.4 | 227 ± 19.8 | 221 ± 14.2 | 118 ± 10.1 | 185 ± 17.2 |
| Male | Hch | 6 | - | 206 ± 18.5 | 239 ± 16.9 | 248 ± 8.8 | 276 ± 20.3 | 299 ± 12.3 | 264 ± 19.3 | 49 ± 6.1 | 275 ± 19.6 | 321 ± 14.1 | 289 ± 22.9 | 237 ± 16 | 144 ± 15.3 | 238 ± 33.5 |
| | 2 mo | 7 | - | 173 ± 20.6 | 201 ± 18.4 | 215 ± 11.1 | 245 ± 14.4 | 318 ± 21.9 | 312 ± 21.7 | 51 ± 9.1 | 315 ± 21.1 | 322 ± 13.5 | 253 ± 16.7 | 225 ± 11.2 | 132 ± 11.1 | 198 ± 8.7 |
| | 4 mo | 6 | - | 163 ± 29.7 | 182 ± 26.1 | 194 ± 23.8 | 229 ± 24.2 | 310 ± 18.1 | 313 ± 18.6 | 43 ± 9.5 | 320 ± 22.1 | 324 ± 23.6 | 254 ± 27.4 | 232 ± 14.2 | 119 ± 11.2 | 196 ± 21.7 |
| | 6 mo | 5 | - | 166 ± 12.9 | 192 ± 20.8 | 214 ± 23.2 | 244 ± 25.3 | 335 ± 27.8 | 345 ± 34 | 47 ± 13.9 | 350 ± 28 | 340 ± 28.9 | 241 ± 25.4 | 247 ± 12.2 | 135 ± 3.1 | 209 ± 17 |
| | Adt | 5 | - | 159 ± 11.4 | 192 ± 10.8 | 193 ± 9.3 | 212 ± 8.1 | 328 ± 23.7 | 338 ± 16.6 | 55 ± 5.9 | 338 ± 21.9 | 328 ± 18.5 | 222 ± 10.9 | 218 ± 3.5 | 143 ± 14.3 | 189 ± 11.1 |

**Supplemental Table 3**

Retinal measurements corresponding to the segmentation methodology applied to obtain global cell populations across the retinal sections. Outer retinal lengths in µm are reported for bins N5-N1, CF, T1-T2, TF, and T4 per sex and age group. Retinal depths in µm were calculated by measuring the depth of the retina at each bin designation. N5 was not assigned a retinal depth because the marginal zone was not measured and the segmentation separating N5 and N4 was designated the retinal depth at N4. N1.5 and T1.5 represent the regions within N1 and T1 that had the largest retinal depth. Pit Center between the CF and T1 bins represents the pit depth as measured by the distance from the retinal pigmented epithelium to the beginning of the center foveal pit. Pit Center between the TF and T4 bins represents the pit depth as measured by the distance from the retinal pigmented epithelium to the beginning of the temporal foveal pit.

### Retina Nuclear Layer Cell Counts

#### GCL Cell Counts (Mean ± SD)

| SEX | AGE | n | N5 | N4 | N3 | N2 | N1 | CF | T1 | T2 | TF | T3 | Total |
|---|---|---|---|---|---|---|---|---|---|---|---|---|---|
| Female | Hch | 5 | 68 ± 20 | 139 ± 26 | 172 ± 26 | 186 ± 22 | 228 ± 47 | 175 ± 68 | 219 ± 35 | 148 ± 13 | 150 ± 19 | 68 ± 11 | 1625 ± 102 |
| | 2 mo | 6 | 80 ± 22 | 120 ± 34 | 151 ± 32 | 160 ± 28 | 245 ± 55 | 253 ± 76 | 220 ± 69 | 133 ± 21 | 148 ± 28 | 67 ± 19 | 1645 ± 255 |
| | 4 mo | 4 | 68 ± 17 | 77 ± 10 | 75 ± 8 | 93 ± 23 | 135 ± 26 | 182 ± 48 | 122 ± 33 | 87 ± 19 | 105 ± 18 | 52 ± 8 | 1041 ± 180 |
| | 6 mo | 6 | 59 ± 14 | 76 ± 11 | 78 ± 8 | 87 ± 16 | 129 ± 30 | 192 ± 47 | 128 ± 24 | 90 ± 14 | 97 ± 25 | 39 ± 15 | 1018 ± 138 |
| | Adt | 5 | 63 ± 10 | 73 ± 8 | 78 ± 15 | 78 ± 17 | 107 ± 30 | 189 ± 65 | 108 ± 25 | 74 ± 8 | 89 ± 23 | 44 ± 9 | 942 ± 150 |
| Male | Hch | 6 | 68 ± 13 | 150 ± 25 | 181 ± 21 | 185 ± 24 | 217 ± 25 | 150 ± 82 | 232 ± 42 | 170 ± 40 | 147 ± 40 | 75 ± 22 | 1659 ± 260 |
| | 2 mo | 7 | 61 ± 19 | 77 ± 18 | 87 ± 24 | 105 ± 29 | 160 ± 46 | 204 ± 89 | 147 ± 49 | 90 ± 31 | 95 ± 31 | 49 ± 10 | 1125 ± 307 |
| | 4 mo | 6 | 76 ± 10 | 82 ± 15 | 86 ± 20 | 96 ± 23 | 140 ± 39 | 222 ± 69 | 131 ± 41 | 86 ± 19 | 110 ± 34 | 58 ± 12 | 1135 ± 203 |
| | 6 mo | 5 | 61 ± 12 | 77 ± 9 | 85 ± 11 | 84 ± 18 | 115 ± 32 | 164 ± 54 | 115 ± 25 | 87 ± 17 | 99 ± 27 | 47 ± 12 | 978 ± 137 |
| | Adt | 4 | 69 ± 9 | 78 ± 14 | 78 ± 9 | 73 ± 8 | 101 ± 24 | 184 ± 63 | 97 ± 19 | 75 ± 7 | 91 ± 21 | 46 ± 9 | 931 ± 143 |

#### INL Cell Counts (Mean ± SD)

| SEX | AGE | n | N5 | N4 | N3 | N2 | N1 | CF | T1 | T2 | TF | T3 | Total |
|---|---|---|---|---|---|---|---|---|---|---|---|---|---|
| Female | Hch | 5 | 549 ± 89 | 1171 ± 95 | 1388 ± 104 | 1521 ± 91 | 1565 ± 195 | 479 ± 337 | 1685 ± 157 | 1365 ± 150 | 856 ± 152 | 634 ± 68 | 11860 ± 707 |
| | 2 mo | 6 | 490 ± 113 | 865 ± 185 | 1072 ± 214 | 1239 ± 220 | 1616 ± 190 | 708 ± 371 | 1595 ± 256 | 1032 ± 152 | 819 ± 152 | 417 ± 137 | 10392 ± 1395 |
| | 4 mo | 4 | 418 ± 76 | 660 ± 67 | 774 ± 121 | 963 ± 97 | 1432 ± 143 | 702 ± 134 | 1442 ± 131 | 893 ± 116 | 760 ± 121 | 360 ± 56 | 8868 ± 741 |
| | 6 mo | 6 | 409 ± 108 | 661 ± 119 | 774 ± 128 | 978 ± 177 | 1491 ± 148 | 977 ± 547 | 1506 ± 156 | 977 ± 164 | 795 ± 101 | 324 ± 99 | 9368 ± 1424 |
| | Adt | 5 | 388 ± 89 | 528 ± 102 | 654 ± 134 | 773 ± 139 | 1275 ± 219 | 885 ± 400 | 1190 ± 189 | 772 ± 79 | 678 ± 108 | 287 ± 79 | 7841 ± 983 |
| Male | Hch | 6 | 549 ± 105 | 1157 ± 123 | 1371 ± 141 | 1488 ± 151 | 1563 ± 131 | 500 ± 445 | 1674 ± 150 | 1375 ± 93 | 853 ± 205 | 638 ± 87 | 11834 ± 1159 |
| | 2 mo | 7 | 456 ± 147 | 754 ± 238 | 883 ± 280 | 1067 ± 361 | 1422 ± 367 | 733 ± 298 | 1358 ± 383 | 925 ± 222 | 752 ± 132 | 421 ± 142 | 9247 ± 2462 |
| | 4 mo | 6 | 392 ± 102 | 658 ± 147 | 804 ± 169 | 1008 ± 196 | 1459 ± 291 | 912 ± 461 | 1389 ± 304 | 951 ± 177 | 822 ± 159 | 374 ± 81 | 9275 ± 1689 |
| | 6 mo | 5 | 347 ± 100 | 552 ± 116 | 705 ± 147 | 887 ± 203 | 1305 ± 361 | 1054 ± 479 | 1290 ± 208 | 869 ± 131 | 756 ± 168 | 339 ± 115 | 8566 ± 1457 |
| | Adt | 4 | 338 ± 62 | 457 ± 78 | 553 ± 91 | 676 ± 128 | 1165 ± 227 | 906 ± 311 | 1118 ± 182 | 699 ± 107 | 655 ± 98 | 258 ± 76 | 7197 ± 1084 |

#### ONL Cell Counts (Mean ± SD)

| SEX | AGE | n | N5 | N4 | N3 | N2 | N1 | CF | T1 | T2 | TF | T3 | Total |
|---|---|---|---|---|---|---|---|---|---|---|---|---|---|
| Female | Hch | 5 | 94 ± 18 | 190 ± 24 | 227 ± 36 | 258 ± 35 | 258 ± 33 | 290 ± 138 | 301 ± 48 | 262 ± 48 | 231 ± 51 | 115 ± 28 | 2350 ± 305 |
| | 2 mo | 6 | 76 ± 24 | 136 ± 43 | 158 ± 39 | 209 ± 63 | 269 ± 51 | 340 ± 88 | 287 ± 67 | 208 ± 46 | 197 ± 52 | 85 ± 37 | 2073 ± 454 |
| | 4 mo | 4 | 66 ± 8 | 92 ± 13 | 97 ± 23 | 127 ± 15 | 211 ± 29 | 256 ± 52 | 224 ± 28 | 141 ± 21 | 157 ± 30 | 53 ± 18 | 1495 ± 118 |
| | 6 mo | 6 | 67 ± 13 | 97 ± 15 | 117 ± 16 | 161 ± 21 | 250 ± 29 | 291 ± 52 | 263 ± 35 | 182 ± 32 | 186 ± 39 | 58 ± 17 | 1755 ± 186 |
| | Adt | 5 | 73 ± 11 | 87 ± 15 | 103 ± 18 | 137 ± 42 | 227 ± 46 | 262 ± 51 | 245 ± 34 | 152 ± 19 | 156 ± 41 | 60 ± 29 | 1579 ± 158 |
| Male | Hch | 6 | 75 ± 18 | 189 ± 38 | 229 ± 38 | 252 ± 30 | 256 ± 39 | 264 ± 54 | 316 ± 45 | 267 ± 32 | 245 ± 35 | 114 ± 23 | 2331 ± 262 |
| | 2 mo | 7 | 69 ± 21 | 107 ± 28 | 131 ± 36 | 178 ± 41 | 226 ± 42 | 282 ± 68 | 244 ± 49 | 177 ± 36 | 179 ± 47 | 68 ± 21 | 1748 ± 340 |
| | 4 mo | 6 | 76 ± 21 | 105 ± 21 | 119 ± 24 | 145 ± 30 | 252 ± 44 | 319 ± 63 | 246 ± 51 | 182 ± 31 | 198 ± 43 | 78 ± 20 | 1816 ± 263 |
| | 6 mo | 5 | 62 ± 21 | 97 ± 23 | 110 ± 28 | 146 ± 38 | 230 ± 53 | 295 ± 74 | 250 ± 54 | 167 ± 29 | 177 ± 60 | 63 ± 22 | 1680 ± 308 |
| | Adt | 4 | 64 ± 17 | 80 ± 13 | 89 ± 10 | 114 ± 28 | 218 ± 60 | 285 ± 44 | 223 ± 48 | 148 ± 24 | 159 ± 37 | 51 ± 10 | 1508 ± 226 |

**Supplemental Table 4**

Mean GCL, INL, and ONL cell counts separated by bin number, sex, and age group. Bins N5-N1, CF, T1-T2, TF, and T4 are denoted along with total cell counts per retinal section.

### Central and Temporal fovea photoreceptor cell counts

**Central ONL Cell Counts (Mean ± SD)**

| SEX | AGE | n | Bin Length (μm) | -20 | -19 | -18 | -17 | -16 | -15 | -14 | -13 | -12 | -11 | -10 | -9 | -8 | -7 | -6 | -5 | -4 | -3 | -2 | -1 |
|---|---|---|---|---|---|---|---|---|---|---|---|---|---|---|---|---|---|---|---|---|---|---|---|
| Female | Hch | 5 | 32 ± 1 | 14 ± 2.2 | 13 ± 2.2 | 14 ± 2.1 | 15 ± 1.9 | 16 ± 1.4 | 17 ± 2.4 | 16 ± 2.2 | 19 ± 2.7 | 18 ± 4.3 | 21 ± 1.3 | 18 ± 2.6 | 20 ± 3.2 | 25 ± 5 | 23 ± 4 | 22 ± 4.3 | 21 ± 2.9 | 13 ± 1.8 | 7 ± 2.1 | 4 ± 1.5 | 0 ± 0 |
| | 2mo | 6 | 43 ± 3.5 | 14 ± 4.3 | 13 ± 3.4 | 14 ± 5.2 | 14 ± 5.2 | 15 ± 4.1 | 15 ± 4.6 | 16 ± 4.2 | 20 ± 7.1 | 20 ± 6.1 | 20 ± 6.9 | 25 ± 8.6 | 26 ± 6 | 29 ± 5.8 | 32 ± 7.7 | 38 ± 10.8 | 35 ± 13.4 | 29 ± 10.1 | 19 ± 6.9 | 10 ± 4 | 1 ± 0.8 |
| | 4mo | 4 | 50 ± 2.6 | 16 ± 3.9 | 15 ± 4.9 | 17 ± 2.4 | 16 ± 2.2 | 16 ± 3.1 | 16 ± 2.2 | 16 ± 2.9 | 16 ± 2.4 | 16 ± 3.6 | 19 ± 4.7 | 21 ± 1.7 | 24 ± 3.8 | 27 ± 4.5 | 31 ± 2.1 | 40 ± 4.9 | 46 ± 5.9 | 44 ± 5.5 | 33 ± 4.9 | 17 ± 4.3 | 3 ± 1 |
| | 6mo | 6 | 53 ± 2.1 | 14 ± 2.5 | 14 ± 2.7 | 14 ± 3.4 | 14 ± 2.8 | 14 ± 3.4 | 14 ± 3 | 14 ± 3.5 | 17 ± 4 | 16 ± 2.3 | 17 ± 2.7 | 18 ± 2.9 | 22 ± 2.9 | 26 ± 5.3 | 30 ± 5 | 33 ± 7.5 | 35 ± 6.2 | 32 ± 8.5 | 25 ± 7.8 | 13 ± 2.6 | 1 ± 1.3 |
| | Adt | 5 | 59 ± 3.2 | 15 ± 1.3 | 15 ± 2.1 | 17 ± 4.2 | 15 ± 2.9 | 17 ± 3.7 | 16 ± 5 | 18 ± 4.2 | 18 ± 1.8 | 17 ± 3.9 | 19 ± 4.8 | 21 ± 7.9 | 21 ± 4.5 | 23 ± 3.2 | 27 ± 7.6 | 35 ± 5.5 | 44 ± 6.3 | 43 ± 4.9 | 31 ± 8 | 15 ± 4.7 | 3 ± 1.5 |
| Male | Hch | 6 | 33 ± 1.1 | 16 ± 4.1 | 15 ± 3.7 | 14 ± 3 | 15 ± 3.3 | 14 ± 3.7 | 17 ± 4.3 | 18 ± 4.6 | 19 ± 6.2 | 20 ± 6.4 | 23 ± 5.4 | 23 ± 4.4 | 23 ± 6.3 | 24 ± 6.8 | 29 ± 7.1 | 27 ± 6.7 | 21 ± 5.3 | 14 ± 3.7 | 9 ± 3.5 | 4 ± 2.1 | 0 ± 0 |
| | 2mo | 7 | 48 ± 2.8 | 17 ± 4.4 | 17 ± 5.5 | 17 ± 5.4 | 17 ± 5.3 | 18 ± 4.1 | 16 ± 5.2 | 17 ± 5.1 | 19 ± 7.1 | 21 ± 6.5 | 22 ± 8 | 25 ± 8.3 | 27 ± 9.1 | 32 ± 10.3 | 36 ± 11.5 | 42 ± 14.6 | 43 ± 12.7 | 38 ± 9.9 | 28 ± 9.2 | 15 ± 5.6 | 2 ± 2 |
| | 4mo | 6 | 57 ± 5.1 | 19 ± 1.7 | 19 ± 2.6 | 19 ± 2.7 | 20 ± 3.2 | 19 ± 2.8 | 19 ± 2.6 | 19 ± 2 | 21 ± 2.2 | 20 ± 2.8 | 21 ± 4.3 | 22 ± 4.7 | 23 ± 5.8 | 27 ± 8.4 | 36 ± 4.8 | 44 ± 6.7 | 48 ± 10 | 54 ± 15 | 43 ± 12.1 | 23 ± 6.9 | 5 ± 4.3 |
| | 6mo | 5 | 62 ± 4.8 | 12 ± 2 | 14 ± 2.5 | 14 ± 4.2 | 14 ± 3.1 | 16 ± 2.7 | 16 ± 3.4 | 14 ± 4.6 | 15 ± 3.8 | 16 ± 6.3 | 16 ± 2.4 | 20 ± 3.8 | 24 ± 5 | 25 ± 1.1 | 27 ± 6.1 | 37 ± 3.6 | 43 ± 7.3 | 48 ± 12.7 | 36 ± 13.4 | 20 ± 7.5 | 2 ± 1.6 |
| | Adt | 4 | 70 ± 4 | 16 ± 3 | 16 ± 3.4 | 17 ± 2.7 | 16 ± 3.8 | 16 ± 3.9 | 17 ± 5.2 | 18 ± 4.1 | 20 ± 3.7 | 21 ± 1 | 20 ± 4.3 | 22 ± 4.3 | 24 ± 4.6 | 28 ± 7.8 | 34 ± 10.7 | 40 ± 10.8 | 47 ± 13.3 | 53 ± 7.3 | 41 ± 7.8 | 27 ± 4.1 | 4 ± 1.9 |

| SEX | AGE | n | CF Pit | 1 | 2 | 3 | 4 | 5 | 6 | 7 | 8 | 9 | 10 | 11 | 12 | 13 | 14 | 15 | 16 | 17 | 18 | 19 | 20 |
|---|---|---|---|---|---|---|---|---|---|---|---|---|---|---|---|---|---|---|---|---|---|---|---|
| Female | Hch | 5 | - | 0 ± 0 | 5 ± 1.9 | 6 ± 3 | 14 ± 1.2 | 18 ± 5.7 | 25 ± 3.3 | 21 ± 4.3 | 23 ± 2.3 | 19 ± 3.1 | 20 ± 5.5 | 20 ± 4.8 | 19 ± 3.4 | 18 ± 3.3 | 16 ± 2.6 | 14 ± 2.9 | 14 ± 3.6 | 12 ± 1.3 | 14 ± 3.8 | 13 ± 1.8 | 13 ± 4.2 |
| | 2mo | 6 | - | 1 ± 0.8 | 10 ± 4.5 | 18 ± 8 | 27 ± 13.2 | 33 ± 10.7 | 37 ± 8.3 | 36 ± 9.7 | 31 ± 7.8 | 25 ± 5.8 | 21 ± 4.2 | 20 ± 4.6 | 18 ± 3.4 | 16 ± 3.4 | 15 ± 4.9 | 13 ± 3 | 14 ± 5 | 14 ± 5.6 | 13 ± 3.3 | 13 ± 2.6 | 13 ± 5.2 |
| | 4mo | 4 | - | 4 ± 1.3 | 19 ± 4.8 | 32 ± 8.4 | 42 ± 8.4 | 47 ± 10.6 | 35 ± 9.7 | 33 ± 3.3 | 25 ± 6.4 | 25 ± 5.3 | 20 ± 2.2 | 19 ± 0.8 | 16 ± 2.2 | 17 ± 2.6 | 17 ± 1.3 | 17 ± 2.6 | 17 ± 4.1 | 17 ± 5.6 | 17 ± 1.2 | 15 ± 1.5 | 16 ± 3.1 |
| | 6mo | 6 | - | 1 ± 1.3 | 13 ± 2.8 | 23 ± 3 | 33 ± 6.8 | 36 ± 6.9 | 33 ± 8 | 28 ± 6.9 | 22 ± 4.8 | 20 ± 4.8 | 16 ± 3.3 | 15 ± 4.1 | 15 ± 2.9 | 13 ± 2.7 | 12 ± 4.5 | 11 ± 2.7 | 11 ± 3.1 | 12 ± 4.2 | 11 ± 2.9 | 10 ± 1.9 | 10 ± 1.8 |
| | Adt | 5 | - | 3 ± 2.1 | 18 ± 4.5 | 36 ± 4.8 | 52 ± 8.5 | 48 ± 7.5 | 32 ± 7.9 | 26 ± 5.6 | 25 ± 7.1 | 22 ± 4.5 | 19 ± 3.8 | 18 ± 5.2 | 15 ± 3.5 | 15 ± 3.8 | 15 ± 2.3 | 15 ± 3.6 | 14 ± 2.8 | 15 ± 2.7 | 16 ± 5.9 | 17 ± 3.7 | 17 ± 3.8 |
| Male | Hch | 6 | - | 0 ± 0 | 5 ± 2.2 | 9 ± 2.9 | 17 ± 4.3 | 22 ± 4 | 25 ± 4.1 | 27 ± 7.3 | 25 ± 4.4 | 24 ± 6.3 | 24 ± 6.5 | 22 ± 5 | 22 ± 5.4 | 19 ± 6 | 18 ± 4 | 16 ± 3.9 | 15 ± 4.6 | 14 ± 2.8 | 14 ± 2.6 | 14 ± 2.4 | 15 ± 3.2 |
| | 2mo | 7 | - | 2 ± 2.2 | 14 ± 5.2 | 25 ± 7 | 37 ± 6.5 | 44 ± 9.3 | 41 ± 11.2 | 34 ± 9.4 | 32 ± 9.1 | 26 ± 10 | 23 ± 8 | 20 ± 7.4 | 18 ± 7.1 | 16 ± 6.3 | 16 ± 5.8 | 13 ± 3.6 | 15 ± 5.3 | 15 ± 4.5 | 15 ± 6.4 | 15 ± 5.3 | 14 ± 4.7 |
| | 4mo | 6 | - | 5 ± 3 | 21 ± 8 | 41 ± 11.2 | 55 ± 13.5 | 50 ± 16.5 | 42 ± 8.7 | 34 ± 8.1 | 27 ± 6.4 | 23 ± 5.6 | 21 ± 3.4 | 19 ± 2.6 | 21 ± 3.9 | 19 ± 1.9 | 17 ± 3.3 | 19 ± 2.9 | 20 ± 3.6 | 19 ± 3.2 | 17 ± 3.8 | 16 ± 2.8 | 17 ± 2.2 |
| | 6mo | 5 | - | 5 ± 1.5 | 19 ± 6.6 | 39 ± 14.7 | 43 ± 11.6 | 45 ± 7.1 | 32 ± 4.3 | 29 ± 5.5 | 20 ± 3 | 20 ± 2.9 | 18 ± 2.4 | 17 ± 4.5 | 17 ± 4.6 | 15 ± 3.5 | 14 ± 2.7 | 15 ± 3.3 | 14 ± 3.9 | 14 ± 2.9 | 14 ± 4.2 | 14 ± 4.4 | 14 ± 5.5 |
| | Adt | 4 | - | 5 ± 1 | 22 ± 4.2 | 46 ± 6.6 | 53 ± 9.7 | 45 ± 9 | 32 ± 9.8 | 26 ± 7.1 | 24 ± 5.4 | 22 ± 3.3 | 20 ± 2.2 | 18 ± 2.6 | 18 ± 2.6 | 19 ± 3 | 18 ± 2.4 | 19 ± 3.7 | 16 ± 2.1 | 15 ± 2.3 | 15 ± 4.3 | 13 ± 3.4 | 12 ± 2.6 |

**Temporal ONL Cell Counts (Mean ± SD)**

| SEX | AGE | n | Bin Length (μm) | -10 | -9 | -8 | -7 | -6 | -5 | -4 | -3 | -2 | -1 | 1 | 2 | 3 | 4 | 5 | 6 | 7 | 8 | 9 | 10 |
|---|---|---|---|---|---|---|---|---|---|---|---|---|---|---|---|---|---|---|---|---|---|---|---|
| Female | Hch | 5 | 38 ± 1.5 | 17 ± 4.7 | 17 ± 3.8 | 18 ± 4.3 | 16 ± 2.6 | 17 ± 4.5 | 18 ± 5 | 20 ± 4.5 | 21 ± 5.9 | 20 ± 6.8 | 20 ± 1.9 | 19 ± 3.1 | 18 ± 4.4 | 18 ± 4.4 | 14 ± 2.8 | 12 ± 4.5 | 9 ± 3.2 | 9 ± 3.1 | 9 ± 2.5 | 7 ± 0.9 | 7 ± 0.8 |
| | 2mo | 6 | 54 ± 4.2 | 18 ± 4 | 18 ± 4.6 | 18 ± 3.7 | 18 ± 4.8 | 19 ± 5.2 | 19 ± 5.2 | 21 ± 2.4 | 25 ± 3.6 | 26 ± 4.5 | 21 ± 5.9 | 22 ± 6.8 | 20 ± 2.8 | 15 ± 2.9 | 13 ± 4 | 11 ± 2.7 | 10 ± 1.5 | 9 ± 1.6 | 8 ± 1.7 | 8 ± 1.4 | 7 ± 2.2 |
| | 4mo | 4 | 64 ± 2.8 | 19 ± 4 | 18 ± 5.5 | 18 ± 2.6 | 17 ± 3.4 | 17 ± 4.3 | 15 ± 2.6 | 16 ± 3.6 | 20 ± 4.5 | 23 ± 5.6 | 24 ± 2.5 | 20 ± 4 | 19 ± 1.3 | 16 ± 5 | 11 ± 1.3 | 11 ± 2.9 | 11 ± 1.3 | 9 ± 2.5 | 8 ± 0.8 | 7 ± 2.6 | 6 ± 1 |
| | 6mo | 6 | 64 ± 2.9 | 18 ± 3.8 | 18 ± 4.2 | 19 ± 5.2 | 18 ± 2.8 | 19 ± 1.9 | 17 ± 3.6 | 20 ± 1.5 | 22 ± 4.9 | 26 ± 5 | 22 ± 4.8 | 21 ± 3.3 | 16 ± 2.3 | 14 ± 3.1 | 11 ± 1.1 | 10 ± 1.5 | 9 ± 1.8 | 9 ± 1.2 | 8 ± 0.8 | 7 ± 0.6 | 6 ± 1.7 |
| | Adt | 5 | 71 ± 3.5 | 18 ± 2.6 | 19 ± 7.2 | 19 ± 5.9 | 20 ± 8.2 | 20 ± 7.4 | 19 ± 8.9 | 21 ± 7.2 | 22 ± 6.5 | 28 ± 5.3 | 29 ± 4.8 | 28 ± 6.6 | 21 ± 6 | 16 ± 4.9 | 13 ± 4.3 | 10 ± 3.8 | 8 ± 0.8 | 7 ± 2.3 | 7 ± 2.2 | 6 ± 1.1 | 6 ± 1.1 |
| Male | Hch | 6 | 39 ± 1.1 | 18 ± 5.3 | 17 ± 5.6 | 18 ± 4.9 | 19 ± 2.9 | 19 ± 3.9 | 20 ± 4.4 | 20 ± 5 | 24 ± 4.7 | 25 ± 9.7 | 23 ± 5.9 | 21 ± 4.6 | 22 ± 7.9 | 19 ± 9.4 | 20 ± 10.3 | 16 ± 7.5 | 14 ± 6.5 | 12 ± 6.6 | 12 ± 6.8 | 11 ± 7 | 12 ± 6.1 |
| | 2mo | 7 | 60 ± 4.6 | 18 ± 5.8 | 16 ± 4.9 | 16 ± 4.6 | 15 ± 3.8 | 16 ± 5.6 | 20 ± 3.9 | 23 ± 6.3 | 23 ± 3.7 | 27 ± 3.8 | 24 ± 6.4 | 22 ± 5.1 | 22 ± 5.7 | 17 ± 4.1 | 14 ± 3.1 | 11 ± 2.4 | 10 ± 1.2 | 9 ± 0.8 | 9 ± 0.8 | 8 ± 1 | 8 ± 1.1 |
| | 4mo | 6 | 67 ± 6.3 | 16 ± 3.4 | 16 ± 1.8 | 15 ± 2.1 | 15 ± 2.9 | 17 ± 2.5 | 17 ± 2.4 | 18 ± 2.6 | 21 ± 3.1 | 25 ± 4.3 | 23 ± 9.5 | 21 ± 6.4 | 17 ± 4.2 | 14 ± 3.4 | 12 ± 2.7 | 10 ± 1 | 10 ± 1 | 9 ± 0.8 | 8 ± 0.4 | 8 ± 1 | 7 ± 0.9 |
| | 6mo | 5 | 76 ± 5.7 | 15 ± 5.2 | 16 ± 4.8 | 16 ± 6.7 | 17 ± 5.8 | 17 ± 3.4 | 20 ± 3.7 | 19 ± 3 | 22 ± 4.1 | 21 ± 3.7 | 25 ± 2.5 | 25 ± 4.7 | 21 ± 5.6 | 15 ± 3.2 | 12 ± 1.6 | 11 ± 1.1 | 10 ± 1.2 | 8 ± 0.8 | 8 ± 1.2 | 7 ± 1.8 | 6 ± 2.9 |
| | Adt | 4 | 83 ± 2.7 | 16 ± 1.4 | 16 ± 2.5 | 16 ± 1 | 16 ± 1 | 15 ± 2.2 | 16 ± 1.8 | 17 ± 2.7 | 20 ± 2.1 | 24 ± 2.6 | 25 ± 4.2 | 27 ± 2.6 | 18 ± 1.7 | 13 ± 3.3 | 11 ± 1.9 | 12 ± 1 | 11 ± 1.5 | 10 ± 1.6 | 8 ± 3.3 | 5 ± 0 | 6 ± 2.1 |

**Supplemental Table 5**

Central and temporal fovea and associated retinal region photoreceptor cell counts. Central fovea cell distribution was measured by mean ONL cell counts within 20 degrees on either side of the center foveal pit (-20 degrees to +20 degrees). The negative degrees represent the optical degrees on the retina extending on the nasal side of the center foveal pit while the positive degrees indicate the optical degrees on the retina extending on the temporal side of the center foveal pit, closer to the temporal fovea. Cell counts are reported per age and per sex, and retinal bin length is reported before cell counts. The break in the table for the central ONL cell counts represents the transition from negative to positive degrees with CF Pit denoted. Temporal fovea cell distribution was measured by mean ONL cell counts within the 10 degrees on either side of the temporal foveal pit. The negative degrees represent the intrafoveal region, which is the retina between the central and temporal fovea pits. The positive degrees indicate the marginal retina, which is the retina on the other side of the temporal fovea pit that is closer to the ciliary marginal zone and cornea. Cell counts are reported per age and per sex, and retinal bin length is reported before cell counts.

#### Central INL Cell Counts (Mean ± SD)

| SEX | AGE | n | Bin Length (µm) | -20 | -19 | -18 | -17 | -16 | -15 | -14 | -13 | -12 | -11 | -10 | -9 | -8 | -7 | -6 | -5 | -4 | -3 | -2 | -1 |
|---|---|---|---|---|---|---|---|---|---|---|---|---|---|---|---|---|---|---|---|---|---|---|---|
| Female | Hch | 5 | 32 ± 1 | 121 ± 16.4 | 122 ± 18 | 124 ± 21.3 | 127 ± 25.5 | 130 ± 22.5 | 139 ± 24.3 | 144 ± 21.3 | 148 ± 16 | 149 ± 19.9 | 141 ± 20.1 | 125 ± 20.2 | 115 ± 16.6 | 100 ± 12.6 | 86 ± 11 | 67 ± 7.5 | 51 ± 7.4 | 39 ± 5 | 21 ± 1.3 | 6 ± 2.1 | 0 ± 0 |
| | 2mo | 6 | 43 ± 3.5 | 153 ± 6.4 | 155 ± 17.5 | 155 ± 12.3 | 159 ± 21.4 | 166 ± 15.5 | 173 ± 11.7 | 183 ± 18.9 | 196 ± 17.1 | 210 ± 18.3 | 213 ± 21.2 | 204 ± 18.1 | 197 ± 30.4 | 185 ± 28.4 | 172 ± 30.1 | 142 ± 26.1 | 112 ± 28.9 | 77 ± 21.7 | 53 ± 22.8 | 20 ± 10.2 | 0 ± 0 |
| | 4mo | 4 | 50 ± 2.6 | 136 ± 13 | 128 ± 15.9 | 138 ± 19.6 | 146 ± 19.3 | 153 ± 23.2 | 152 ± 29.5 | 156 ± 30.5 | 170 ± 27.7 | 185 ± 29.1 | 204 ± 32.8 | 226 ± 44.4 | 226 ± 36.5 | 213 ± 30.2 | 196 ± 23.4 | 188 ± 25.7 | 156 ± 28.9 | 124 ± 33.4 | 80 ± 27 | 34 ± 12.9 | 2 ± 1.7 |
| | 6mo | 6 | 53 ± 2.1 | 119 ± 14.7 | 122 ± 14.5 | 130 ± 19.2 | 135 ± 25.5 | 139 ± 23.7 | 141 ± 14.7 | 144 ± 16.4 | 157 ± 16.7 | 176 ± 25.3 | 183 ± 27.4 | 199 ± 32.6 | 198 ± 24.6 | 189 ± 19.5 | 178 ± 15.6 | 154 ± 18.8 | 128 ± 11.8 | 85 ± 11.4 | 52 ± 8.3 | 22 ± 8.8 | 0 ± 0.5 |
| | Adt | 5 | 59 ± 3.2 | 132 ± 26.8 | 129 ± 17.2 | 133 ± 13.2 | 144 ± 15.7 | 149 ± 20.5 | 148 ± 13.7 | 155 ± 15.5 | 166 ± 20.3 | 171 ± 15.2 | 194 ± 25.6 | 207 ± 19 | 217 ± 25.1 | 229 ± 32.2 | 230 ± 38.9 | 211 ± 39.6 | 167 ± 36.4 | 124 ± 24.3 | 74 ± 13.9 | 32 ± 11.9 | 1 ± 1.3 |
| Male | Hch | 6 | 33 ± 1.1 | 138 ± 20.7 | 140 ± 20.2 | 140 ± 18 | 142 ± 22 | 149 ± 23.6 | 157 ± 24.4 | 163 ± 23.5 | 166 ± 22.9 | 156 ± 25.2 | 153 ± 27 | 139 ± 20.4 | 127 ± 16 | 117 ± 10.3 | 102 ± 15.3 | 78 ± 12.8 | 58 ± 10.6 | 41 ± 7.4 | 23 ± 3.1 | 5 ± 3.1 | 0 ± 0 |
| | 2mo | 7 | 48 ± 2.8 | 161 ± 34.4 | 167 ± 33 | 172 ± 35.5 | 173 ± 28.2 | 171 ± 24.9 | 176 ± 20.4 | 184 ± 25.9 | 202 ± 25.1 | 210 ± 37.9 | 214 ± 41.1 | 222 ± 40.1 | 219 ± 33.4 | 211 ± 33.1 | 198 ± 30.5 | 174 ± 36 | 145 ± 33 | 106 ± 29.1 | 66 ± 30.9 | 29 ± 19 | 1 ± 2.2 |
| | 4mo | 6 | 57 ± 5.1 | 153 ± 17.9 | 159 ± 20.5 | 162 ± 18 | 166 ± 16.9 | 172 ± 19.6 | 178 ± 20.6 | 180 ± 17.3 | 189 ± 21.3 | 192 ± 20.6 | 213 ± 25.2 | 230 ± 26.1 | 239 ± 25.8 | 240 ± 34.9 | 241 ± 36.4 | 223 ± 31 | 191 ± 40.4 | 150 ± 28.5 | 101 ± 25.8 | 51 ± 15.4 | 6 ± 3.7 |
| | 6mo | 5 | 62 ± 4.8 | 124 ± 27.1 | 128 ± 28.8 | 125 ± 25.5 | 129 ± 29.4 | 133 ± 30.8 | 143 ± 33.9 | 147 ± 33.9 | 152 ± 36 | 160 ± 37.4 | 180 ± 20.8 | 199 ± 28.7 | 217 ± 35.1 | 221 ± 36.9 | 213 ± 29.2 | 211 ± 18 | 186 ± 19.2 | 146 ± 10.6 | 95 ± 13.7 | 42 ± 10 | 3 ± 2.4 |
| | Adt | 4 | 70 ± 4 | 128 ± 11.5 | 142 ± 6.1 | 134 ± 11.4 | 147 ± 4.8 | 148 ± 6.9 | 155 ± 10 | 170 ± 9.7 | 183 ± 15.4 | 190 ± 15.1 | 211 ± 7.5 | 221 ± 12.5 | 234 ± 15.6 | 252 ± 16.7 | 263 ± 23.1 | 258 ± 28.2 | 233 ± 48.8 | 182 ± 45.9 | 107 ± 35.3 | 51 ± 29.7 | 2 ± 2.2 |

| SEX | AGE | n | CF Pit | 1 | 2 | 3 | 4 | 5 | 6 | 7 | 8 | 9 | 10 | 11 | 12 | 13 | 14 | 15 | 16 | 17 | 18 | 19 | 20 |
|---|---|---|---|---|---|---|---|---|---|---|---|---|---|---|---|---|---|---|---|---|---|---|---|
| Female | Hch | 5 | - | 0 ± 0 | 6 ± 1.6 | 22 ± 3.2 | 38 ± 3.5 | 53 ± 6.1 | 71 ± 6.7 | 86 ± 11.3 | 109 ± 13.7 | 125 ± 10.5 | 132 ± 16.9 | 146 ± 15.8 | 161 ± 13.4 | 167 ± 18.1 | 165 ± 28.4 | 146 ± 29.3 | 126 ± 28 | 114 ± 23.9 | 116 ± 22.6 | 112 ± 21.5 | 116 ± 22.2 |
| | 2mo | 6 | - | 0 ± 0.4 | 22 ± 10.2 | 54 ± 18.7 | 82 ± 23.1 | 111 ± 28.4 | 145 ± 30.4 | 175 ± 29.7 | 196 ± 33.3 | 216 ± 28.7 | 231 ± 22.9 | 238 ± 24.3 | 227 ± 10.5 | 199 ± 46.7 | 172 ± 40.4 | 140 ± 29.1 | 138 ± 21.7 | 142 ± 16.6 | 145 ± 7.4 | 138 ± 5.4 | 131 ± 6.5 |
| | 4mo | 4 | - | 2 ± 2.9 | 38 ± 16.4 | 78 ± 32 | 120 ± 34.2 | 158 ± 28.5 | 184 ± 23 | 214 ± 22.9 | 228 ± 19 | 241 ± 29.6 | 256 ± 20.5 | 248 ± 28.5 | 206 ± 30.3 | 170 ± 25.1 | 153 ± 8.4 | 146 ± 23.8 | 141 ± 16.9 | 137 ± 24 | 132 ± 19.1 | 132 ± 18.2 | 125 ± 14.8 |
| | 6mo | 6 | - | 1 ± 1 | 21 ± 5.2 | 59 ± 8.5 | 102 ± 13.3 | 139 ± 11.9 | 171 ± 10.2 | 184 ± 12.7 | 192 ± 17.5 | 199 ± 26.6 | 196 ± 27 | 189 ± 30.1 | 170 ± 19.9 | 135 ± 17.9 | 122 ± 13.5 | 125 ± 18.6 | 124 ± 20.3 | 116 ± 15.9 | 108 ± 12.8 | 108 ± 13.3 | 102 ± 12.3 |
| | Adt | 5 | - | 2 ± 1.7 | 38 ± 12.5 | 82 ± 15.8 | 128 ± 20.6 | 171 ± 14.6 | 202 ± 13.3 | 222 ± 19.9 | 228 ± 29.4 | 220 ± 29.4 | 210 ± 24.3 | 182 ± 23.6 | 159 ± 22.3 | 145 ± 14.1 | 142 ± 5.1 | 134 ± 8.8 | 121 ± 5.7 | 117 ± 13.8 | 125 ± 10 | 124 ± 6.9 | 125 ± 7.5 |
| Male | Hch | 6 | - | 0 ± 0 | 7 ± 3.2 | 26 ± 5.1 | 43 ± 8.4 | 67 ± 10.6 | 86 ± 12.9 | 102 ± 13 | 120 ± 19.4 | 140 ± 16 | 147 ± 23.1 | 159 ± 27.1 | 169 ± 29 | 175 ± 28.1 | 172 ± 32.1 | 155 ± 36.4 | 141 ± 42.4 | 134 ± 27.1 | 128 ± 19.4 | 130 ± 11.7 | 129 ± 13.7 |
| | 2mo | 7 | - | 2 ± 2.1 | 30 ± 19.2 | 68 ± 26.3 | 110 ± 32.4 | 149 ± 31.5 | 187 ± 27.7 | 207 ± 34.8 | 219 ± 39.6 | 232 ± 37.3 | 225 ± 36.6 | 224 ± 47.5 | 220 ± 53.1 | 191 ± 38.2 | 151 ± 24.1 | 143 ± 23.1 | 150 ± 26.2 | 144 ± 25.5 | 144 ± 26.9 | 139 ± 25.6 | 136 ± 18.8 |
| | 4mo | 6 | - | 5 ± 2.8 | 53 ± 15 | 107 ± 22.2 | 161 ± 31 | 202 ± 31.9 | 233 ± 28.9 | 251 ± 32.2 | 262 ± 33.4 | 254 ± 28.4 | 230 ± 38.3 | 208 ± 46.4 | 177 ± 30.2 | 166 ± 18.9 | 164 ± 22 | 161 ± 22.2 | 153 ± 25 | 153 ± 24.9 | 148 ± 24.7 | 153 ± 23 | 139 ± 19.8 |
| | 6mo | 5 | - | 4 ± 2.3 | 46 ± 7.4 | 97 ± 15.8 | 149 ± 20.2 | 187 ± 21.7 | 209 ± 22.2 | 221 ± 35 | 224 ± 44.4 | 209 ± 48.3 | 194 ± 49.5 | 164 ± 37.9 | 145 ± 30.9 | 141 ± 29 | 130 ± 25.6 | 123 ± 25.7 | 118 ± 24.9 | 119 ± 21.9 | 114 ± 17.7 | 112 ± 18.4 | 111 ± 15.5 |
| | Adt | 4 | - | 4 ± 3.1 | 54 ± 25.1 | 115 ± 28.9 | 194 ± 35.4 | 230 ± 25.2 | 249 ± 19.5 | 253 ± 8.8 | 240 ± 19.6 | 226 ± 25.2 | 202 ± 34.3 | 188 ± 18.1 | 166 ± 17.9 | 160 ± 20.2 | 145 ± 18.5 | 139 ± 18.3 | 135 ± 20 | 126 ± 22.9 | 126 ± 18.6 | 126 ± 22.7 | 119 ± 27.2 |

#### Temporal INL Cell Counts (Mean ± SD)

| SEX | AGE | n | Bin Length (µm) | -10 | -9 | -8 | -7 | -6 | -5 | -4 | -3 | -2 | -1 | 1 | 2 | 3 | 4 | 5 | 6 | 7 | 8 | 9 | 10 |
|---|---|---|---|---|---|---|---|---|---|---|---|---|---|---|---|---|---|---|---|---|---|---|---|
| Female | Hch | 5 | 38 ± 1.5 | 133 ± 42.2 | 132 ± 38 | 121 ± 33.7 | 119 ± 31.3 | 124 ± 33.4 | 127 ± 28.3 | 116 ± 27.5 | 98 ± 22.6 | 73 ± 15.5 | 46 ± 9.2 | 50 ± 9.4 | 73 ± 20.2 | 87 ± 17.4 | 98 ± 21.7 | 99 ± 23.5 | 92 ± 24.4 | 90 ± 20.7 | 79 ± 18.4 | 70 ± 19.7 | 59 ±17.8 |
| | 2mo | 6 | 54 ± 4.2 | 182 ± 26.6 | 172 ± 29 | 163 ± 20.4 | 163 ± 19.5 | 168 ± 21 | 175 ± 35 | 171 ± 26.1 | 160 ± 18.2 | 127 ± 15.4 | 69 ± 8.5 | 61 ± 11.4 | 114 ± 16.2 | 129 ± 14.9 | 126 ± 13 | 112 ± 14.4 | 97 ± 12.8 | 88 ± 13.6 | 79 ± 9.7 | 72 ± 11.5 | 63 ± 17.4 |
| | 4mo | 4 | 64 ± 2.8 | 158 ± 30.5 | 145 ± 30.9 | 141 ± 21.7 | 141 ± 17.7 | 142 ± 15.2 | 141 ± 11.9 | 144 ± 16.7 | 145 ± 16.9 | 122 ± 31.9 | 74 ± 20.5 | 70 ± 12.7 | 112 ± 15.8 | 126 ± 23.9 | 115 ± 17.4 | 100 ± 15.6 | 88 ± 12.3 | 75 ± 9.6 | 61 ± 12 | 52 ± 10.6 | 41 ± 11.7 |
| | 6mo | 6 | 64 ± 2.9 | 167 ± 45.4 | 157 ± 38.2 | 154 ± 41 | 155 ± 33.7 | 150 ± 24.8 | 158 ± 24.1 | 163 ± 30.3 | 160 ± 33 | 128 ± 26.3 | 66 ± 21.1 | 65 ± 21.2 | 120 ± 21 | 132 ± 19.4 | 123 ± 17.5 | 107 ± 16.5 | 94 ± 15.8 | 82 ± 12 | 71 ± 8.4 | 61 ± 8.2 | 52 ± 10.2 |
| | Adt | 5 | 71 ± 3.5 | 151 ± 11.6 | 143 ± 13.8 | 146 ± 14 | 145 ± 17.2 | 150 ± 18.8 | 158 ± 22 | 158 ± 17.5 | 152 ± 15.8 | 127 ± 24.5 | 80 ± 23.5 | 71 ± 21 | 117 ± 29.4 | 126 ± 24.5 | 125 ± 30.5 | 102 ± 18.8 | 87 ± 15.9 | 72 ± 9.6 | 62 ± 6.1 | 52 ± 10 | 35 ± 11.6 |
| Male | Hch | 6 | 39 ± 1.1 | 147 ± 40.2 | 142 ± 33.5 | 141 ± 26.2 | 146 ± 25.1 | 147 ± 28.9 | 142 ± 25 | 132 ± 20.8 | 112 ± 19.2 | 83 ± 14.5 | 57 ± 13.5 | 58 ± 10.5 | 83 ± 13.5 | 110 ± 10.7 | 124 ± 17.9 | 124 ± 22.6 | 122 ± 19.1 | 108 ± 20.6 | 105 ± 19 | 95 ± 26.8 | 90 ± 35.6 |
| | 2mo | 7 | 60 ± 4.6 | 164 ± 24.2 | 156 ± 26.8 | 155 ± 29.6 | 150 ± 25.4 | 156 ± 28 | 161 ± 26.2 | 167 ± 31.1 | 154 ± 25.4 | 125 ± 24 | 73 ± 18.5 | 65 ± 16.6 | 116 ± 25.1 | 136 ± 25.9 | 126 ± 23 | 114 ± 18.5 | 103 ± 13.7 | 91 ± 11.2 | 79 ± 10.7 | 75 ± 9.1 | 59 ± 8.8 |
| | 4mo | 6 | 67 ± 6.3 | 163 ± 26.2 | 162 ± 28.7 | 153 ± 28.4 | 149 ± 32.2 | 149 ± 25.3 | 156 ± 28.6 | 174 ± 27.4 | 168 ± 23.9 | 135 ± 22.8 | 77 ± 26.6 | 70 ± 28.4 | 121 ± 20.2 | 136 ± 22 | 127 ± 19.8 | 114 ± 20.7 | 98 ± 19 | 83 ± 13.1 | 70 ± 13.4 | 58 ± 9.2 | 49 ± 10.9 |
| | 6mo | 5 | 76 ± 5.7 | 188 ± 38.7 | 178 ± 32.2 | 173 ± 35.6 | 176 ± 33.6 | 175 ± 23.4 | 179 ± 26.7 | 187 ± 19.7 | 185 ± 26.7 | 155 ± 30.1 | 91 ± 20.7 | 86 ± 13.5 | 145 ± 16.3 | 159 ± 15 | 134 ± 13.7 | 119 ± 15.4 | 96 ± 12.1 | 86 ± 15.8 | 74 ± 12.1 | 59 ± 17.5 | 42 ± 24.1 |
| | Adt | 4 | 83 ± 2.7 | 151 ± 28.3 | 142 ± 18.7 | 140 ± 24.7 | 141 ± 20.5 | 142 ± 15.3 | 144 ± 9 | 157 ± 15.1 | 154 ± 18.3 | 142 ± 21.3 | 89 ± 26.8 | 84 ± 21.5 | 131 ± 16.8 | 132 ± 15 | 118 ± 8.3 | 89 ± 14.1 | 72 ± 10 | 60 ± 6.3 | 54 ± 10 | 43 ± 2.1 | 40 ± 1 |

**Supplemental Table 6**

Central and temporal fovea and associated retinal region INL cell counts. For the central fovea, the negative degrees represent the optical degrees on the retina extending on the nasal side of the center foveal pit while the positive degrees indicate the optical degrees on the retina extending on the temporal side of the center foveal pit, closer to the temporal fovea. Cell counts are reported per age and per sex, and retinal bin length is reported before cell counts. The break in the table for the central INL cell counts represents the transition from negative to positive degrees with CF Pit denoted. For the temporal fovea, the negative degrees represent the intrafoveal region, which is the retina between the central and temporal fovea pits. The positive degrees indicate the marginal retina, which is the retina on the other side of the temporal fovea pit that is closer to the ciliary marginal zone and cornea. Cell counts are reported per age and per sex, and retinal bin length is reported before cell counts.

##### Central GCL Cell Counts (Mean ± SD)

| SEX | AGE | n | Bin Length (µm) | -20 | -19 | -18 | -17 | -16 | -15 | -14 | -13 | -12 | -11 | -10 | -9 | -8 | -7 | -6 | -5 | -4 | -3 | -2 | -1 |
|---|---|---|---|---|---|---|---|---|---|---|---|---|---|---|---|---|---|---|---|---|---|---|---|
| Female | Hch | 5 | 32 ± 1 | 20 ± 4.3 | 23 ± 3.8 | 23 ± 4.7 | 22 ± 5.8 | 24 ± 3.8 | 22 ± 3.3 | 20 ± 1.1 | 17 ± 2.3 | 18 ± 3.4 | 19 ± 4.7 | 19 ± 3.3 | 20 ± 2.1 | 21 ± 5.3 | 24 ± 3.1 | 24 ± 2.4 | 25 ± 4.7 | 25 ± 4.2 | 24 ± 5.5 | 13 ± 4 | 0 ± 0 |
| | 2mo | 6 | 43 ± 3.5 | 20 ± 4.1 | 21 ± 6.1 | 23 ± 6.3 | 25 ± 6.7 | 27 ± 5.3 | 29 ± 4.6 | 28 ± 3.2 | 28 ± 6.4 | 26 ± 4.3 | 25 ± 2.4 | 23 ± 2.3 | 23 ± 1 | 23 ± 2.8 | 24 ± 3.5 | 28 ± 6.4 | 31 ± 7.1 | 35 ± 10.1 | 42 ± 11.5 | 37 ± 7.4 | 3 ± 3.4 |
| | 4mo | 4 | 50 ± 2.6 | 18 ± 2.1 | 19 ± 3 | 18 ± 4.4 | 21 ± 3.9 | 20 ± 2.8 | 23 ± 2.4 | 27 ± 7.3 | 28 ± 6.7 | 28 ± 3.1 | 29 ± 2.9 | 27 ± 7 | 28 ± 5 | 30 ± 2.2 | 31 ± 3.7 | 31 ± 3.8 | 31 ± 2.9 | 35 ± 4.5 | 40 ± 11.5 | 42 ± 15.6 | 11 ± 12.3 |
| | 6mo | 6 | 53 ± 2.1 | 17 ± 3.3 | 19 ± 5.5 | 17 ± 3.7 | 20 ± 3.9 | 18 ± 2.7 | 24 ± 4.1 | 28 ± 6.6 | 26 ± 5.4 | 27 ± 4.4 | 30 ± 4 | 29 ± 4.5 | 27 ± 1.9 | 28 ± 3.9 | 31 ± 4.8 | 30 ± 3.9 | 29 ± 5.4 | 35 ± 5.2 | 37 ± 5.7 | 28 ± 11.6 | 3 ± 3.5 |
| | Adt | 5 | 59 ± 3.2 | 17 ± 4.3 | 17 ± 1.8 | 17 ± 2.3 | 19 ± 0.8 | 22 ± 4.9 | 25 ± 6.8 | 28 ± 7.8 | 29 ± 5.2 | 28 ± 1.9 | 35 ± 8 | 36 ± 6.7 | 33 ± 9.1 | 33 ± 4.6 | 33 ± 8.9 | 33 ± 6.8 | 33 ± 7 | 33 ± 4.6 | 37 ± 3.8 | 35 ± 9.5 | 9 ± 9.3 |
| Male | Hch | 6 | 33 ± 1.1 | 24 ± 1.9 | 25 ± 4.8 | 28 ± 4.6 | 27 ± 3.4 | 28 ± 5.9 | 28 ± 11.1 | 23 ± 3.9 | 22 ± 3.8 | 20 ± 4.2 | 21 ± 2.3 | 22 ± 2.8 | 22 ± 3.9 | 25 ± 4.2 | 25 ± 4.2 | 27 ± 5.8 | 30 ± 5.2 | 31 ± 5.8 | 28 ± 7.7 | 16 ± 9 | 0 ± 0.4 |
| | 2mo | 7 | 48 ± 2.8 | 20 ± 3.1 | 22 ± 5.5 | 24 ± 3.8 | 21 ± 3.3 | 25 ± 5.4 | 26 ± 6.7 | 29 ± 7.3 | 27 ± 6.6 | 28 ± 5.8 | 26 ± 5.8 | 28 ± 8.5 | 28 ± 6.5 | 27 ± 5.9 | 29 ± 6.8 | 31 ± 6.3 | 30 ± 6.1 | 35 ± 7.7 | 39 ± 11.2 | 39 ± 19.6 | 12 ± 23.1 |
| | 4mo | 6 | 57 ± 5.1 | 16 ± 2.6 | 18 ± 2.4 | 19 ± 2 | 18 ± 2.4 | 21 ± 3.7 | 23 ± 3.3 | 23 ± 3.5 | 27 ± 5 | 28 ± 3.4 | 32 ± 5 | 31 ± 5.5 | 29 ± 4.4 | 29 ± 5.6 | 28 ± 6.3 | 30 ± 6.5 | 34 ± 5.9 | 35 ± 5.5 | 41 ± 7.2 | 45 ± 7.6 | 16 ± 10.2 |
| | 6mo | 5 | 62 ± 4.8 | 17 ± 5.5 | 16 ± 4.5 | 15 ± 4.1 | 17 ± 3.7 | 19 ± 4.7 | 20 ± 5.9 | 20 ± 3.5 | 21 ± 4 | 24 ± 6.2 | 29 ± 5.2 | 29 ± 6.1 | 28 ± 2 | 27 ± 3.8 | 30 ± 3.2 | 29 ± 4.6 | 33 ± 4.7 | 35 ± 7.2 | 39 ± 5.4 | 41 ± 6.2 | 8 ± 8.6 |
| | Adt | 4 | 70 ± 4 | 16 ± 2.6 | 16 ± 2.2 | 15 ± 2.1 | 16 ± 1 | 17 ± 2.3 | 19 ± 2.6 | 22 ± 1.4 | 24 ± 2.4 | 28 ± 3.8 | 33 ± 9.1 | 35 ± 7.9 | 37 ± 5.4 | 36 ± 6.9 | 33 ± 3.5 | 33 ± 5 | 33 ± 5.2 | 35 ± 1.3 | 39 ± 2.1 | 39 ± 9 | 6 ± 8 |

| SEX | AGE | n | CF Pit | 1 | 2 | 3 | 4 | 5 | 6 | 7 | 8 | 9 | 10 | 11 | 12 | 13 | 14 | 15 | 16 | 17 | 18 | 19 | 20 |
|---|---|---|---|---|---|---|---|---|---|---|---|---|---|---|---|---|---|---|---|---|---|---|---|
| Female | Hch | 5 | - | 0 ± 0 | 14 ± 4.7 | 23 ± 3.7 | 27 ± 5.4 | 27 ± 3.6 | 27 ± 2.3 | 26 ± 1.8 | 23 ± 3.4 | 22 ± 3.8 | 20 ± 4.4 | 21 ± 3.2 | 18 ± 3.4 | 19 ± 3.4 | 19 ± 2.2 | 24 ± 5.5 | 28 ± 4.8 | 30 ± 12.1 | 22 ± 4.9 | 23 ± 5.5 | 23 ± 5.9 |
| | 2mo | 6 | - | 4 ± 4.6 | 37 ± 4.2 | 41 ± 5.5 | 35 ± 3.7 | 31 ± 5.2 | 29 ± 2.5 | 27 ± 3.8 | 25 ± 3.7 | 25 ± 5.3 | 24 ± 4.2 | 23 ± 3.2 | 26 ± 4.5 | 32 ± 6.6 | 36 ± 13.8 | 36 ± 20.6 | 30 ± 12.9 | 23 ± 5.2 | 23 ± 5.4 | 22 ± 6.6 | 21 ± 7.5 |
| | 4mo | 4 | - | 11 ± 16.3 | 44 ± 19.4 | 39 ± 9.2 | 39 ± 9.9 | 33 ± 7.3 | 32 ± 8.3 | 32 ± 1 | 29 ± 2.4 | 28 ± 1.7 | 28 ± 2.2 | 30 ± 2.5 | 38 ± 12.8 | 43 ± 14.8 | 35 ± 13.1 | 29 ± 6.9 | 23 ± 6 | 19 ± 2.8 | 22 ± 3.8 | 19 ± 3.6 | 23 ± 4.5 |
| | 6mo | 6 | - | 2 ± 2 | 34 ± 12 | 41 ± 7.1 | 38 ± 8.2 | 34 ± 5.4 | 32 ± 3.6 | 31 ± 3.6 | 29 ± 3.4 | 30 ± 4.9 | 26 ± 3.2 | 26 ± 3.1 | 31 ± 5.3 | 36 ± 15.2 | 26 ± 4.4 | 24 ± 6.2 | 21 ± 4.3 | 18 ± 3.9 | 18 ± 3 | 18 ± 3.3 | 16 ± 2.9 |
| | Adt | 5 | - | 7 ± 5 | 42 ± 9.5 | 37 ± 2.2 | 36 ± 4 | 35 ± 4.3 | 34 ± 8.7 | 31 ± 7.5 | 34 ± 9.4 | 34 ± 5.1 | 37 ± 8.2 | 37 ± 8.6 | 29 ± 4.9 | 26 ± 4.5 | 23 ± 3 | 23 ± 4.1 | 21 ± 3.2 | 22 ± 3 | 20 ± 3.4 | 21 ± 2.2 | 18 ± 3.3 |
| Male | Hch | 6 | - | 2 ± 2.5 | 20 ± 6 | 30 ± 3.8 | 32 ± 4.4 | 30 ± 5.9 | 30 ± 4.9 | 27 ± 5.6 | 26 ± 6 | 26 ± 5.2 | 22 ± 3.4 | 21 ± 2.8 | 21 ± 3.9 | 20 ± 2.8 | 22 ± 2.6 | 26 ± 5.2 | 30 ± 9.4 | 29 ± 7.2 | 28 ± 4.1 | 26 ± 3.8 | 24 ± 3.4 |
| | 2mo | 7 | - | 11 ± 15.1 | 38 ± 18 | 39 ± 9.2 | 38 ± 10.2 | 34 ± 6.9 | 30 ± 5.9 | 30 ± 5.5 | 30 ± 5.9 | 26 ± 6 | 29 ± 5.1 | 32 ± 4.8 | 28 ± 5.8 | 30 ± 10.1 | 39 ± 21 | 28 ± 8.5 | 25 ± 5 | 22 ± 5.1 | 21 ± 5 | 21 ± 6.4 | 21 ± 6.4 |
| | 4mo | 6 | - | 18 ± 11.6 | 49 ± 15.4 | 43 ± 8.7 | 37 ± 8.8 | 34 ± 5.2 | 31 ± 4.4 | 29 ± 4.3 | 30 ± 3.9 | 32 ± 5.7 | 33 ± 7.7 | 36 ± 7.6 | 30 ± 9.4 | 30 ± 6.9 | 25 ± 4.2 | 25 ± 6.1 | 22 ± 5.6 | 24 ± 4.3 | 24 ± 3.1 | 20 ± 2.3 | 22 ± 3.5 |
| | 6mo | 5 | - | 10 ± 8.2 | 44 ± 7.8 | 38 ± 7.2 | 35 ± 6.8 | 31 ± 4.5 | 27 ± 2.3 | 26 ± 4.3 | 28 ± 1.5 | 33 ± 5.9 | 30 ± 5.3 | 29 ± 5.2 | 30 ± 9.4 | 24 ± 6.1 | 23 ± 4.4 | 19 ±6.1 | 17 ± 3.6 | 17 ± 5.9 | 19 ± 6 | 16 ± 3 | 16 ± 2.6 |
| | Adt | 4 | - | 7 ± 7.9 | 37 ± 9.5 | 39 ± 5.9 | 39 ± 5.4 | 35 ± 11.4 | 33 ± 6.5 | 39 ± 5.1 | 39 ± 5.1 | 33 ± 2.5 | 34 ± 4.1 | 29 ± 7 | 25 ± 3 | 22 ± 4.9 | 22 ± 4.5 | 21 ± 2.8 | 19 ± 1 | 19 ± 3.5 | 19 ± 3.3 | 18 ± 4.9 | 20 ± 8.4 |

##### Temporal GCL Cell Counts (Mean ± SD)

| SEX | AGE | n | Bin Length | -10 | -9 | -8 | -7 | -6 | -5 | -4 | -3 | -2 | -1 | 1 | 2 | 3 | 4 | 5 | 6 | 7 | 8 | 9 | 10 |
|---|---|---|---|---|---|---|---|---|---|---|---|---|---|---|---|---|---|---|---|---|---|---|---|
| Female | Hch | 5 | 38 ± 1.5 | 25 ± 7 | 24 ± 6.3 | 22 ± 6.2 | 22 ± 5 | 22 ± 6.6 | 22 ± 6.4 | 22 ± 5.2 | 24 ± 7.2 | 24 ± 4.3 | 2 ± 4.7 | 25 ± 5.6 | 25 ± 5 | 21 ± 4.7 | 18 ± 3.2 | 17 ± 5.9 | 16 ± 6.7 | 14 ± 3.7 | 14 ± 4 | 12 ± 3.7 | 10 ± 4 |
| | 2mo | 6 | 54 ± 4.2 | 27 ± 4 | 26 ± 6.2 | 26 ± 5.2 | 31 ± 10.2 | 28 ± 7.1 | 28 ± 4.8 | 23 ± 4.3 | 24 ± 3.9 | 28 ± 4.9 | 33 ± 12.7 | 32 ± 13 | 28 ± 6.3 | 21 ± 4.7 | 18 ± 3.3 | 17 ± 3 | 18 ± 5.1 | 16 ± 5 | 16 ± 8.6 | 10 ± 4.8 | 8 ± 3.2 |
| | 4mo | 4 | 64 ± 2.8 | 20 ± 4.7 | 22 ± 4 | 20 ± 4.1 | 23 ± 2.9 | 23 ± 2.9 | 24 ± 5.1 | 26 ± 3.6 | 27 ± 5.4 | 31 ± 5 | 29 ± 8 | 25 ± 5.5 | 24 ± 2.4 | 20 ± 1.8 | 18 ± 3.4 | 14 ± 3.9 | 13 ± 2.6 | 13 ± 3 | 9 ± 2.1 | 8 ± 1.3 | 6 ± 2.1 |
| | 6mo | 6 | 64 ± 2.9 | 24 ± 4.3 | 25 ± 3.8 | 24 ± 3.3 | 22 ± 3.1 | 27 ± 5.7 | 25 ± 5.1 | 26 ± 0.9 | 28 ± 2.4 | 32 ± 4.1 | 34 ± 10 | 33 ± 10.3 | 29 ± 5.8 | 22 ± 2.2 | 18 ± 2 | 17 ± 1.9 | 14 ± 2.1 | 13 ± 2.2 | 11 ± 1.2 | 10 ± 1.7 | 7 ± 2.7 |
| | Adt | 5 | 71 ± 3.5 | 21 ± 2.7 | 21 ± 2.2 | 22 ± 3.4 | 20 ± 2.7 | 22 ± 2.6 | 23 ± 2.3 | 27 ± 2.9 | 26 ± 1.9 | 26 ± 3 | 29 ± 9 | 27 ± 8.2 | 25 ± 2.6 | 22 ± 1.6 | 17 ± 4.4 | 16 ± 6.6 | 12 ± 2.8 | 10 ± 2.1 | 9 ± 2.2 | 7 ± 2.3 | 5 ± 3.9 |
| Male | Hch | 6 | 39 ± 1.1 | 25 ± 8.4 | 25 ± 5.4 | 26 ± 7.6 | 26 ± 5.5 | 25 ± 5.5 | 22 ± 3.3 | 26 ± 4.4 | 27 ± 4.4 | 35 ± 5 | 36 ± 8.9 | 33 ± 5 | 34 ± 6.7 | 28 ± 5.4 | 22 ± 3.1 | 21 ± 3.8 | 20 ± 4.9 | 19 ± 3.9 | 18 ± 4.5 | 15 ± 5.2 | 14 ± 6.9 |
| | 2mo | 7 | 60 ± 4.6 | 24 ± 6.5 | 26 ± 9.6 | 26 ± 6.6 | 27 ± 6 | 28 ± 8.1 | 26 ± 6.5 | 26 ± 6.9 | 29 ± 5.3 | 29 ± 4.6 | 32 ± 9.1 | 31 ± 10.2 | 27 ± 5.4 | 23 ± 4.4 | 19 ± 6.6 | 17 ± 5.3 | 14 ± 5.4 | 15 ± 6.4 | 13 ± 4.1 | 11 ± 3.8 | 9 ± 4.2 |
| | 4mo | 6 | 67 ± 6.3 | 22 ± 3.2 | 22 ± 2.6 | 23 ± 3.8 | 25 ± 2.9 | 26 ± 5.6 | 26 ± 4.1 | 25 ± 3.4 | 28 ± 6.2 | 30 ± 5.5 | 33 ± 6.5 | 25 ± 6.7 | 27 ± 3.1 | 24 ± 5.7 | 19 ± 1.4 | 16 ± 1.6 | 15 ± 3.1 | 14 ± 2.1 | 12 ± 2.5 | 10 ± 1.8 | 8 ± 4.1 |
| | 6mo | 5 | 76 ± 5.7 | 23 ± 4.2 | 25 ± 4 | 27 ± 5.5 | 25 ± 3.7 | 27 ± 7.1 | 26 ± 8.8 | 29 ± 6.3 | 33 ± 5.8 | 34 ± 3.9 | 37 ± 8.9 | 37 ± 8.6 | 33 ± 4.5 | 23 ± 5 | 20 ± 2.9 | 18 ± 3 | 15 ± 2.8 | 15 ± 2.3 | 11 ± 2.3 | 9 ± 3.6 | 6 ± 5.3 |
| | Adt | 4 | 83 ± 2.7 | 25 ± 9.5 | 23 ± 7.9 | 24 ± 5.1 | 22 ± 3.3 | 26 ± 3 | 29 ± 8.3 | 31 ± 5.3 | 34 ± 5.3 | 34 ± 7.8 | 28 ± 5.7 | 27 ± 8.8 | 26 ± 4 | 21 ± 2.2 | 20 ± 1.7 | 14 ± 1.7 | 12 ± 2.9 | 11 ± 0.5 | 8 ± 3.3 | 8 ± 1.2 | 7 ± 1 |

**Supplemental Table 7**

Central and temporal fovea and associated retinal region GCL cell counts. For the central fovea, the negative degrees represent the optical degrees on the retina extending on the nasal side of the center foveal pit while the positive degrees indicate the optical degrees on the retina extending on the temporal side of the center foveal pit, closer to the temporal fovea. Cell counts are reported per age and per sex, and retinal bin length is reported before cell counts. The break in the table for the central GCL cell counts represents the transition from negative to positive degrees with CF Pit denoted. For the temporal fovea, the negative degrees represent the intrafoveal region, which is the retina between the central and temporal fovea pits. The positive degrees indicate the marginal retina, which is the retina on the other side of the temporal fovea pit that is closer to the ciliary marginal zone and cornea. Cell counts are reported per age and per sex, and retinal bin length is reported before cell counts.
